# Activity-assembled nBAF complex mediates rapid immediate early gene transcription by regulating RNA Polymerase II productive elongation

**DOI:** 10.1101/2023.12.30.573688

**Authors:** Karen G. Cornejo, Andie Venegas, Morgan H. Sono, Madeline Door, Brenda Gutierrez-Ruiz, Lucy B. Karabedian, Supratik G. Nandi, Emily C Dykhuizen, Ramendra N Saha

## Abstract

Signal-dependent RNA Polymerase II (Pol2) productive elongation is an integral component of gene transcription, including those of immediate early genes (IEGs) induced by neuronal activity. However, it remains unclear how productively elongating Pol2 overcome nucleosomal barriers. Using RNAi, three degraders, and several small molecule inhibitors, we show that the mammalian SWI/SNF complex of neurons (neuronal BAF, or nBAF) is required for activity-induced transcription of neuronal IEGs, including *Arc*. The nBAF complex facilitates promoter-proximal Pol2 pausing, signal-dependent Pol2 recruitment (loading), and importantly, mediates productive elongation in the gene body via interaction with the elongation complex and elongation-competent Pol2. Mechanistically, Pol2 elongation is mediated by activity-induced nBAF assembly (especially, ARID1A recruitment) and its ATPase activity. Together, our data demonstrate that the nBAF complex regulates several aspects of Pol2 transcription and reveal mechanisms underlying activity-induced Pol2 elongation. These findings may offer insights into human maladies etiologically associated with mutational interdiction of BAF functions.

## Introduction

BRG1/BRM-associated factor (BAF) complexes are conserved ATP-dependent chromatin remodelers. They were first discovered in yeast as the SWI/SNF (SWItch/Sucrose Non-Fermentable) complex^1–3^, then in Drosophila^4^, and in mammals soon after^5,6^. Mammalian BAF complexes are now known to be multimeric large complexes weighing 1-2 MDa, which are formed combinatorially with products of 31 genes. Three biochemically distinct BAF complexes are delineated in mammals: canonical BAF (cBAF), polybromo-associated BAF (PBAF), and GLTSCR1-BAF (GBAF)^7^, also called non-canonical BAF (ncBAF)^8^ (Figure 1a). All three complexes feature either SMARCA4 (BRG1) or SMARCA2 (BRM) as the ATPase core unit, which interacts with several other subunits to form functional complexes. The quintessential cBAF complex utilizes 12-15 subunits, including, SMARCC1 (BAF155) and/or SMARCC2 (BAF170) as scaffolding subunits^9–12^, and ARID1A or ARID1B as mutually exclusive subcomplex-defining subunits. Combinatorial assemblies around these core components in a modular fashion permute several functional BAF complexes^13,14^.

**Figure 1:**
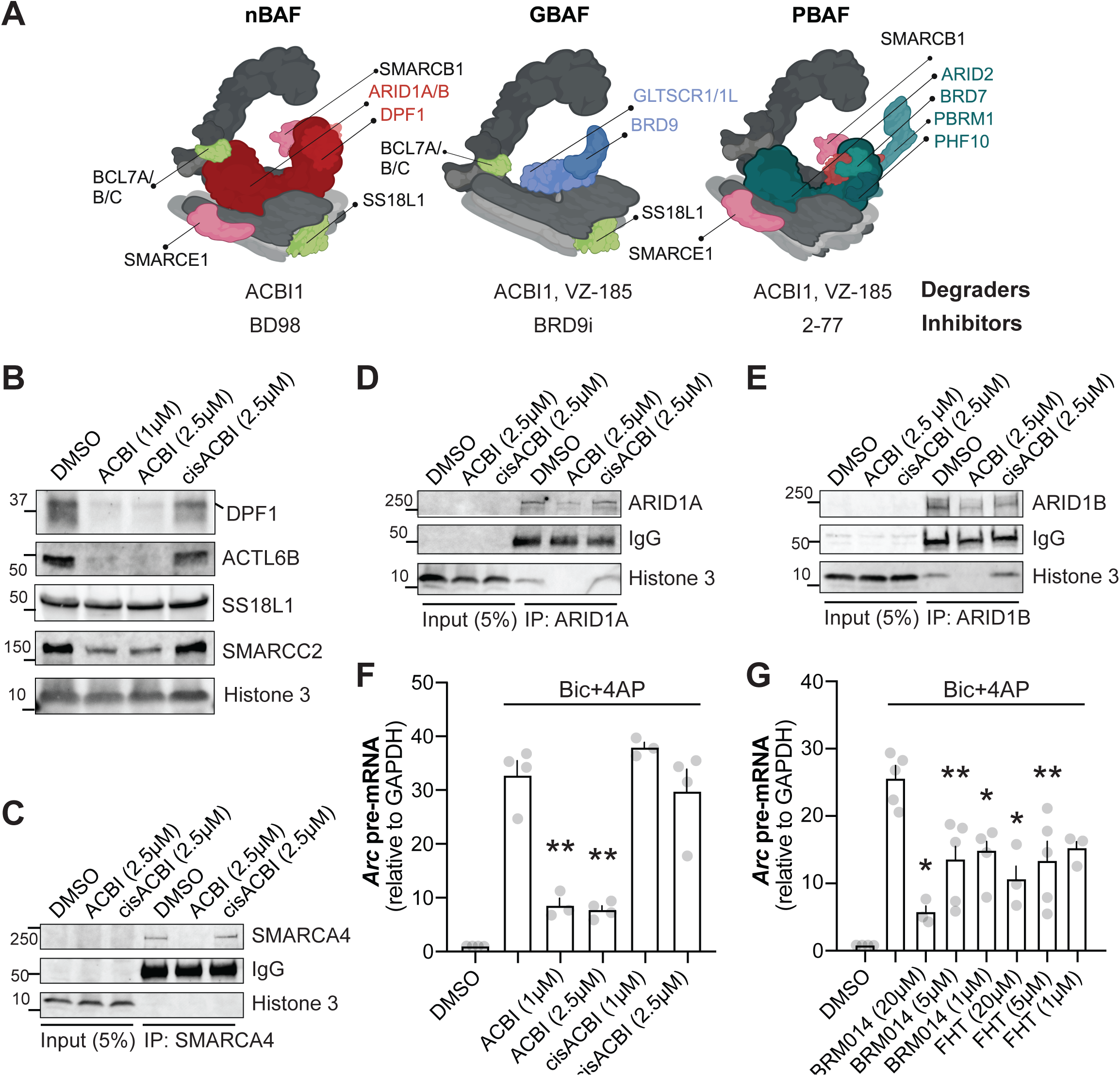
BAF complex is required for optimal *Arc* transcription. A) Schematic representation of three biochemically distinct BAF complexes: nBAF (neuronal cBAF), GBAF, and PBAF. Complex defining unique subunits are represented in colors. Subunits shared by all three complexes are not indicated. Degraders and small molecule inhibitors to target each complex are noted below. Schematic was created using BioRender. B) Neurons were treated with DMSO (control), indicated concentration of ACBI1 or cisACBI (inactive isomer of ACBI1) for 3 hours. Whole-cell lysates were electrophoresed, Western blotted, and probed for indicated BAF subunits. C, D, and E): Neuronal lysates were used to immunoprecipitate SMARCA4 (C), ARID1A (D), and ARID1B (E). 5% of cell lysates was used as input. Histone 3 and IgG are depicted as the loading control. F-G) Transcriptional assays where *Arc* pre-mRNA normalized by GAPDH pre-mRNA levels are illustrated. F) Neurons were treated with ACBI1 or cisACBI for three hours followed by bicuculline and 4AP treatment for 15 minutes (Bic+4AP). G) Neurons treated with BAF ATPase domain inhibitors, BRM014 and FHT for 30 minutes, followed by Bic+4AP treatment to induce neuronal activity. Grey dots represent biological replicates, errors bar show SE of the mean. \**P*<0.05; \*\**P*< 0.01. One-way ANOVA analyses were followed by Tukey’s *post hoc* test. Approximate position of nearest molecular weight marker is depicted against each band.

BAF complexes drive transcriptional programs, such as those mediating differentiation and lineage specification of developing cells^15^. Developmental programs often include compositional transformations of BAF complexes. For example, during neurogenesis, neuronal progenitor version of the cBAF complex (npBAF), which features SS18, DPF2 and ACTL6A (BAF53a), swaps several subunits to become the neuronal BAF (nBAF) complex, which is instead characterized by SS18L1 (CREST), DPF1/3 and ACTL6B (BAF53b)^10^. The transition from npBAF to nBAF also includes shifts in subunit stoichiometry, such as increasing ratios of SMARCC2 compared to SMARCC1. The nBAF is both ‘necessary’ and ‘sufficient’ for neurogenesis. Forcing nBAF expression in human fibroblasts converts them to neurons^16,17^, while knocking out nBAF subunits in developing neurons causes defective synaptogenesis^18,19^. Moreover, mutations in genes encoding several BAF subunits disrupt neurodevelopment^10,12,20–22^ and coincide with human neuro-developmental disorders (NDDs), including intellectual disability (ID) and autism spectrum disorders (ASD)^20,23–27^. A recent study reported that cBAF subunit coding genes possess the greatest number of *de novo* missense and protein-truncating mutations among all nuclear protein complexes^28^. Together, it is clear that nBAF drives neuron-specific gene transcription in developing and mature neurons^19,29^, and NDDs may result from loss of such function. However, underlying molecular mechanisms of such transcriptional processes are understood vaguely.

BAF complexes are primarily alluded to as chromatin remodelers, which refers to their ability to reposition, evict, or disassemble nucleosomes. Related to this quidditative function, BAF complexes also play several roles in gene transcription. In yeast, the SWI/SNF complex is known to actively decondense chromatin, antagonize chromatin-mediated transcriptional repression, and act as gene transcription activators^30,31^. In mammals, BAF complexes work in concert with many transcription factors to promote chromatin accessibility and transcription^32^. They are enriched at promoters and enhancers, where they are implicated in promoter-enhancer interactions, enhancer maintenance, and activation of lineage-specific enhancers^33–36^. Recently, they were correlated with promoter-proximal Pol2 pausing^37^ and shown to be necessary for activation of inducible promoters^38^. However, it remains unclear if BAF complexes are necessary for Pol2 productive elongation.

Productive elongation requires a signal-directed switch of Pol2 from its obligatory promoter proximal paused state to active transcription. This process is orchestrated by positive transcription elongation factor b (P-TEFb), the master regulator of transcription elongation^39^. CDK9, the constituent kinase in P-TEFb, phosphorylates position 2 serines (S2) in C-terminal domain (CTD) heptad repeats of Rpb1, the largest Pol2 subunit. The phosphorylated CTD acts as a scaffold for elongation factors such as SPT6, which assemble to form the elongation complex (EC)^40,41^. The EC increases elongation competency and drives Pol2 several folds faster than isolated Pol2^42^. Notwithstanding such insights however, it remains unclear how Pol2 overcome nucleosomes in the gene body during productive elongation. Elongating Pol2 complexes can surmount nucleosomal barriers without needing remodelers^43^, but inefficiently. Therefore, it is proposed that efficient elongation is aided by nucleosome remodelers. However, the relationship between the EC and any such remodeler remains unclear.

Indirect evidence in the literature suggests that the BAF complex may have a role during Pol2 elongation. SWI/SNF components were reported in transcribing gene bodies of yeast and *Drosophila*^44–49^. Brm immunostaining in *Drosophila* polytene chromatin localizes to active chromatin and depletion of Brm drastically reduced global transcription^46^. In yeast, Swi2 and Pol2 occupancy was identical in the coding region of an osmotically-induced gene^44^. SMARCA2 and SMARCA4 are found in the coding region of several genes, where they aid alternative splicing^47^. Finally, in a cellular artificial construct, Brg1 facilitated nucleosome traversal by Pol2^45^. Taken together, association of the BAF complex with Pol2 elongation is indicated, but rules of such engagement or underlying mechanisms thereof remain obscure, especially in the mammalian system.

To investigate any role of nBAF in Pol2 elongation, we focused on neuronal activity-induced gene transcription. We have previously shown that in response to activity at the membrane, neurons undertake a gene transcription program, whereby three distinct classes of genes are briskly expressed in temporally distinct waves: rapid immediate early genes (rIEGs), delayed immediate early genes (dIEGs), and secondary genes that require *de novo* protein translation^50^. The earliest among them are rIEGs, a small subset of 15-18 genes (e.g., *Arc*) whose transcription happens when few or no other genes undergo inducible transcription. Promoters of rIEGs have open chromatin and are preloaded with necessary transcription factors and paused Pol2 nearby^50,51^. Upon stimulation, the pioneer Pol2 undertakes productive elongation and pre-mRNA production within minutes. Additional rounds of Pol2 recruitment enables a robust transcriptional response. This process requires the fast acting MAP kinase pathway^50^, which regulates several transcription factors and also the positive transcription elongation factor b (pTEFb)^52^. Using rIEG *Arc* as a model gene, we demonstrate in dissociated primary rat neurons that the nBAF mediates activity-induced rapid transcription. We also show that the nBAF complex assembles in response to activity, interacts as a key component with the elongation complex, and utilizes ATPase activity to facilitate Pol2 elongation.

## Results

### BAF complex and its ATPase activity is necessary for activity-induced rIEG transcription

Does activity-induced rapid neuronal gene transcription require the BAF complex? To answer this question, we utilized several pharmacological inhibitors, degraders, and RNAi. First, we used the recently developed ACBI1, the proteolysis targeting chimera (PROTAC) degrader of BAF ATPases (SMARCA2 and SMARCA4) and associated subunits^53^. Dissociated rat cortical neurons were treated with ACBI1 for three hours and several nBAF subunits levels were estimated. ACBI1, but not its negative control (inactive *cis*-ACBI1), reduced DPF1, ACTL6B, and SMARCA4 beneath detection levels (Figure 1B and 1C). Levels of SMARCC2, ARID1A, ARID1B, and SS18L1 were reduced but not eliminated (Figure 1B, 1D, and 1E). Therefore, ACBI1 degrades major BAF complex subunits to various degrees but leaves intact some protein orphans from the complex, such as SMARCC2 and SS18L1. Next, sustained synaptic activity was stimulated with Bicuculline and 4-Aminopyridine (Bic+4AP) for 15 minutes and transcriptional response was assessed by quantifying *Arc* pre-mRNA^54,55^. Fifteen minutes of activity was chosen to isolate transcription of rIEGs from other activity-induced transcription, which are detectable later in the hour^50,51^. ACBI1, but not *cis*-ACBI1, significantly attenuated activity-induced *Arc* transcription (Figure 1F). To rule out an *Arc*-specific effect, eight other rIEGs were additionally tested. All eight rIEGs displayed sensitivity to ACBI1, but not to *cis*-ACBI1 (Figure S1A), suggesting a general role of the complex in activity-induced rIEG transcription. Furthermore, significant transcriptional impairment also resulted when neurons were pre-incubated with two small-molecule allosteric BAF ATPase inhibitors, BRM014 and FHT1015^56,57^ (Figure 1G). Together, the combined use of pharmacological degradation and inhibition of the BAF complex ATPase function suggests that BAF plays a key role in activity-induced transcription of *Arc* and other rIEGs.

To further validate the involvement of BAF complex in activity-induced transcription of IEGs, we knocked down two BAF complex subunits via short hairpin RNA. Dissociated cortical neurons were infected with RNAi targeting SMARCC2^58^ or ARID1A, the scaffolding and nBAF-defining subunits respectively. Protein levels of both targeted proteins were reduced confirming RNAi efficacy (Figure 2A and 2B). Consistent with above assays involving the degrader and ATPase inhibitors, SMARCC2 and ARID1A knockdown significantly impaired *Arc* transcription following Bicuculline treatments (Figure 2C). Adduced together, these data portray the BAF complex as a positive regulator of activity-induced transcription. Also, among three BAF complexes, transcriptional impairment due to ARID1A depletion suggests involvement of the nBAF complex in the process.

**Figure 2.**
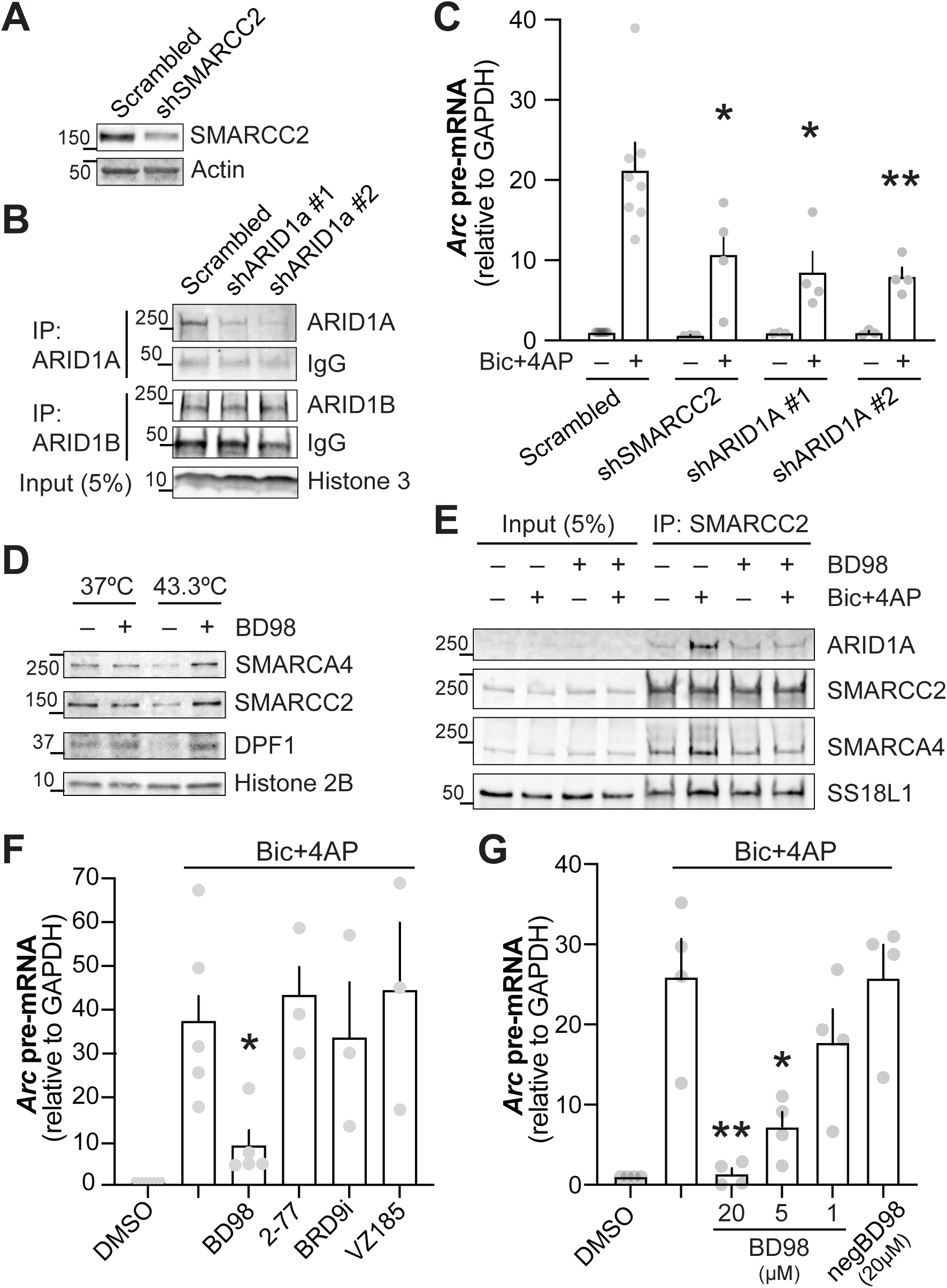
RNAi-dependent depletion and pharmacological perturbation of nBAF complex subunits attenuate *Arc* transcription. A and B) Protein levels indicating knockdown of SMARCC2 and ARID1A. Neurons were infected with lentiviruses to deliver shRNA targeted against *SMARCC2* (three days) or *ARID1A* (five days). Scrambled shRNA was used as control. Knockdown of SMARCC2 (A; whole cell lysate), ARID1A (B; immunoprecipitation of whole cell lysate) was verified by protein levels. Specificity of ARID1A RNAi is depicted by with ARID1B immunoprecipitation, whose levels remained unaffected. C) Neurons depleted of indicated nBAF subunits were treated with bicuculline and 4AP (Bic+4AP; neuronal activity) for 15 minutes. Normalized *Arc* pre-mRNA levels are displayed. D) Cellular thermal shift assay (CETSA) was performed on live cells treated with BD98 (20 µM) for 30 minutes. Cell lysates were then analyzed via Western blot to assess thermal stability of SMARCC2, SMARCA4, and nBAF-specific DPF1. All three BAF subunits displayed stability at higher temperature in BD98 treated neurons indicating direct binding of the inhibitor with nBAF complex. E) Co-IP assay was performed in lysates from neurons pre-incubated with BD98 (5µM) followed by Bic+4AP treatment for 15 minutes. Anti-SMARCC2 immunoprecipitated samples were separated on a gel and blotted for other BAF subunits. 5% of cell lysate was used as input. F) Neurons were pre-incubated for 30 minutes with BD98 (20µM), BRD7i (5µM), and BRD9i (0.2 µM) to inhibit nBAF, PBAF and GBAF respectively, followed by Bic+4AP for 15 minutes. Normalized *Arc* pre-mRNA levels are displayed G) Similar assay as in F, except neurons were preincubated with indicated doses of BD98. Negative BD98 was used as a control. Grey dots represent biological replicates, error bars show SE of the mean. \**P*<0.05; \*\**P*<0.01. One-way ANOVA analyses were performed followed by Tukey’s *post hoc* test. Approximate position of nearest molecular weight marker is depicted against each band.

### Activity-assembled nBAF complex drives rIEG transcription

Several recently described nBAF, PBAF, and GBAF-specific inhibitors were used next to characterize the BAF complex regulating rIEG transcription. First, we used BD98, a new cBAF inhibitor that specifically inhibits ARID1A complexes^59^. To validate direct binding of BD98 with nBAF complex, we performed cellular thermal shift assay (CETSA^60^) that reports thermal stabilization of proteins upon ligand binding in cells. Treating neurons with BD98 enhanced thermal stability of SMARCC2, SMARCA4, and nBAF-specific DPF1 (Figure 2D) suggesting direct binding of BD98 with the nBAF complex. CETSA could not be performed with ARID1A due to its low abundance. Instead, we performed co-immunoprecipitation assays of endogenous proteins (henceforth, Co-IP) in the presence of BD98. The inhibitor was expected to disrupt nBAF subunit interactions, especially those of ARID1A. Here, we made an unexpected discovery. Compared to the untreated control, interactions of SMARCC2 with ARID1A, SMARCA4, and SS18L1 increased after Bic+4AP treatment (Figure 2E), suggesting an activity-induced assembly of the nBAF complex. This assembly — especially recruitment of ARID1A — was inhibited by BD98 (Figure 2E).

Next, the effect of BD98 on rIEG transcription was assessed along with PBAF- and GBAF-specific inhibitors (2-77^61^ and I-BRD9^62^, respectively). Pre-treatment with BD98, but not PBAF or GBAF inhibitors, significantly attenuated activity-induced transcription of *Arc* and other rIEGs (Figure 2F and S1A). Furthermore, we also used VZ185, the BRD7 and BRD9 PROTAC degrader^63^. Three hours of treatment with VZ185 depleted BRD7, but not nBAF subunit DPF1 or the core subunit SMARCC2 (Figure S1B). We did not detect BRD9 in neuronal extracts with two commercially available antibodies. Like PBAF and GBAF inhibitors, and in contrast to BD98, VZ185 did not affect activity-induced *Arc* induction (Figure 2F). BD98 acted in a dose-dependent manner, where 5µM significantly inhibited *Arc* transcription (Figure 2G) and was therefore used as the BD98 dose for all other assays. Towards due diligence, we checked for any non-specific upstream effects of BD98 by testing its impact on activity-induced MAP kinase activation, a key signaling pathway for rIEGs^50^, and found none at 5µM or less concentrations (Figure S1C).

To expand on activity-induced BAF complex assembly, additional Co-IPs were performed. First, an activity time-course and Co-IPs with SMARCC2 and ARID1A revealed that signal-induced nBAF assembly above the basal level occurs within 15 minutes of treatment and starts to decline thereafter (Figure 3A and 3B). Activity-induced assembly of nBAF was strongly sensitive to an inhibitor cocktail for glutamate- and voltage-gated synaptic transmission, but not noticeably to Tetrodotoxin (TTX; Figure 3C), which inhibits neuronal firing but not synaptic activity. Because synaptic activity communicates with the nucleus via Ca^2+^ as the second messenger^64^, we next tested its role in the process by treating neurons with cell permeable Ca^2+^-chelator BAPTA-AM. Chelation of calcium completely attenuated activity-induced nBAF assembly, but not impacting its basal level (Figure 3D).

**Figure 3.**
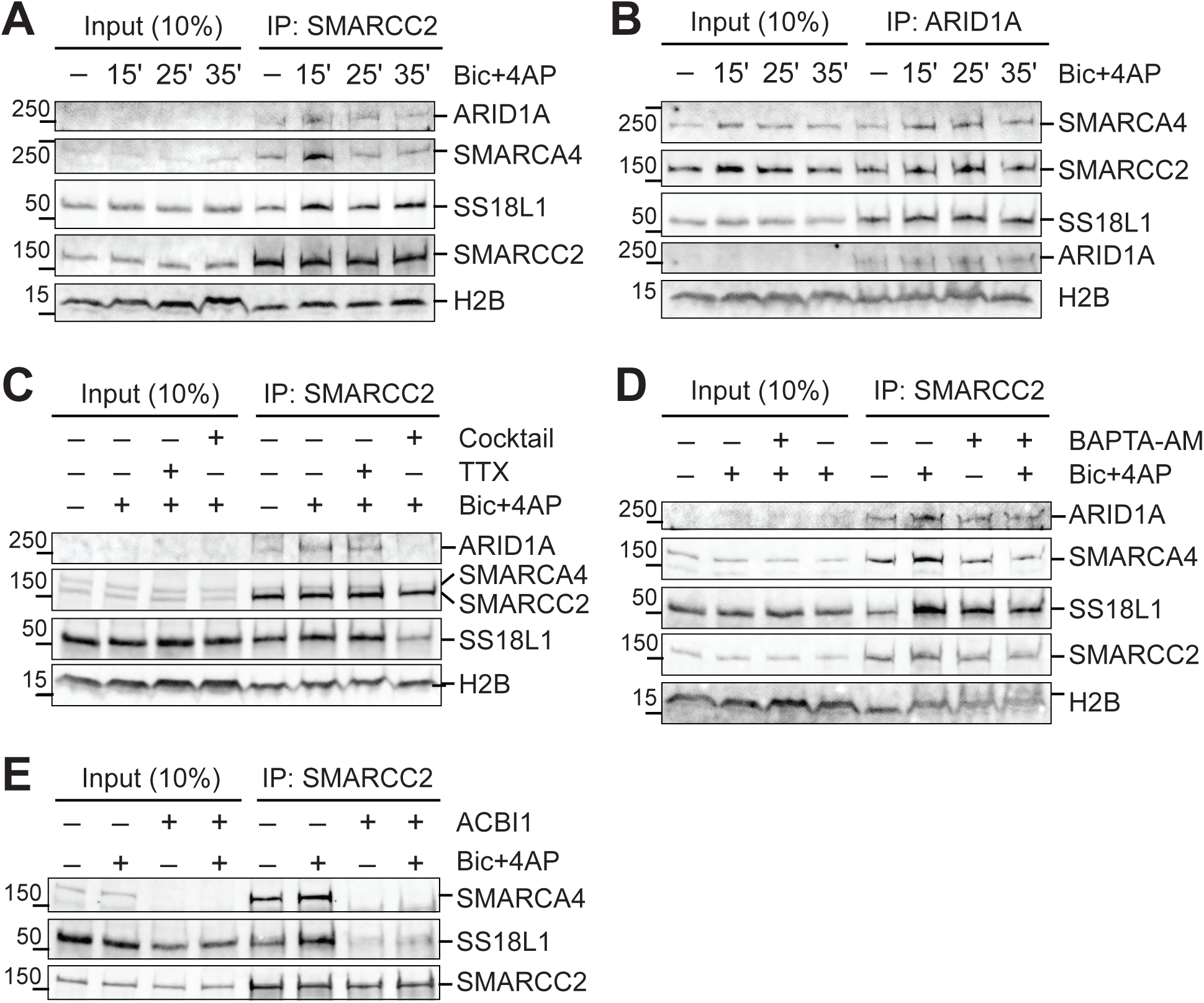
Neuronal activity dynamically assembles nBAF. Co-IP assays with endogenous proteins were performed to detect interactions of BAF subunits in response to activity and various inhibitors or degraders. Immunoprecipitation was performed using anti-SMARCC2 (A, C, D, and E) or anti-ARID1A (B). A-B) Neurons were stimulated with Bic+4AP for indicated time periods. C) Neurons were pre-treated with a cocktail of synaptic inhibitors RS-CPP (2.5 µM), CNQX (2.5 µM) and Nimodipine (5 µM) or TTX (1 µM) to inhibit synaptic activity and neuronal firing respectively, which was followed by 15 minutes of Bic+4AP treatment to induce activity. D) Neurons were incubated with the Ca^2+^ chelator BAPTA-AM (100 µM, 15 minutes) and then treated with Bic+4AP for 15 minutes. E) Neurons were incubated with ACBI1 (2.5 µM) for 3 hours followed by 15 minutes of activity. SMARCA4 degradation confirms ACBI1 efficiency. 10% of cell lysates were used as input. Co-IP material was electrophoresed and Western blotted to detect interacting subunits. Approximate position of nearest molecular weight marker is depicted.

BAF complex assembly has been recently demonstrated to be modular^14^, where the ATPase forms an independent module along with SS18 (in non-neuronal cells) and finalizes complex assembly by binding to the SMARCC-containing core and ARID modules. In figure 2E, hinting at such modular assembly, BD98-mediated inhibition of ARID1A binding to SMARCC2 also prevented activity-induced enhanced binding of SMARCA4. Similarly, degradation of SMARCA4 with ACBI1, which spares both SMARCC2 and SS18L1 to various degrees, inhibited activity-induced enhanced interaction between the two (Figure 3E). This observation suggests that SS18L1 assembles with SMARCC2 only in the presence of the ATPase, again, hinting at modular assembly for the complex.

### Presence of nBAF is necessary for promoter-proximal Pol2 pausing

Prior to investigating the role of nBAF in Pol2 elongation, we first asked if nBAF was relevant for other aspects of the transcription cycle that precede productive elongation. Promoters of rIEGs are characterized by paused Pol2 near their transcription start site, which serves as a mechanism for their rapid induction^51^. The first hint of a synergy between the BAF complex and Pol2 pausing came from our chromatin immunoprecipitation (ChIP) assays where SMARCC2 knockdown attenuated Pol2 enrichments near *Arc* promoter (Figure S2A and S2B). ChIP assays were performed with two antibodies: 1) a monoclonal antibody to the N-terminal region of Rpb1, which detects total Pol2 independent of its phosphorylation states, and 2) a monoclonal antibody to Rpb1 CTD heptads with phosphorylation at serine 5 residues (Rpb1-pS5), which detects promoter bound but elongation incompetent (paused) Pol2^65^. Next, we treated neurons with ACBI or BD98 for 18 hours and assessed Pol2 subunit protein levels. BAF complex degradation or inhibition of nBAF did not alter levels of total Rpb1 or Rpb1-pS5 (Figure S2C). However, consistent with SMARCC2 KD experiments, such treatments depleted paused Pol2 levels near the *Arc* promoter (Figure S2D-S2F). In addition to above-mentioned pair of Rpb1 antibodies, we also used an antibody against Spt5, a transcription factor known to stabilize Pol2 at promoter-proximal regions as part of the pausing complex^66^. Data from all three ChIP assays agreed and suggested that the presence of nBAF is necessary for maintenance of promoter proximal Pol2 pausing. Interestingly however, treatment with ACBI1 or BD98 for shorter durations (up to an hour) did not alter promoter proximal Pol2 levels (data not shown). Put together, the nBAF complex — or certain subunits thereof — facilitate Pol2 pausing, where the facilitation process likely has a low turnover rate.

### Activity-induced Pol2 recruitment and promoter-loading requires BAF subunit SMARCC2

Activity induces paused Pol2 to escape the promoter while also recruiting and promoter-loading several additional rounds of Pol2 to produce a robust transcriptional output^51,67^. Prior to testing its function in elongation, we therefore verified any role of the BAF complex in activity-induced Pol2 recruitment and promoter loading. We performed ChIP assays with antibodies against total Rpb1, Rpb1-pS5, and SMARCC2. SMARCC2 was chosen as it is part of the BAF core module and has DNA-binding domains (SANT and SWIRM) that facilitates cross-linking and immunoprecipitation of chromatin. Neuronal activity significantly enriched Rpb1-pS5, total Pol2, and SMARCC2 levels near *Arc* promoter, which were attenuated in neurons depleted of SMARCC2 (Figure 4B-4D). SMARCC2 signal depletion after its RNAi (Figure 4D) attests to specificity of the anti-SMARCC2 antibody. Furthermore, in agreement with RNAi data, BAF complex degradation with ACBI1 attenuated activity-induced enrichment of Rpb1-pS5 and total Pol2 near *Arc* promoter (Figure 4E and 4F). Unexpectedly however, ACBI1 treatment did not affect promoter proximal SMARCC2 enrichment (Figure 4G). Taken together, this data set indicates that activity-induced Pol2 recruitment is downstream of SMARCC2 enrichment and, as indicated by Figure 4B and 4C, is dependent on it as well. These data also lend support to an activity-dependent modular mode nBAF assembly, where SMARCC2 independently translocates to the promoter region even when several other BAF subunits are degraded.

**Figure 4.**
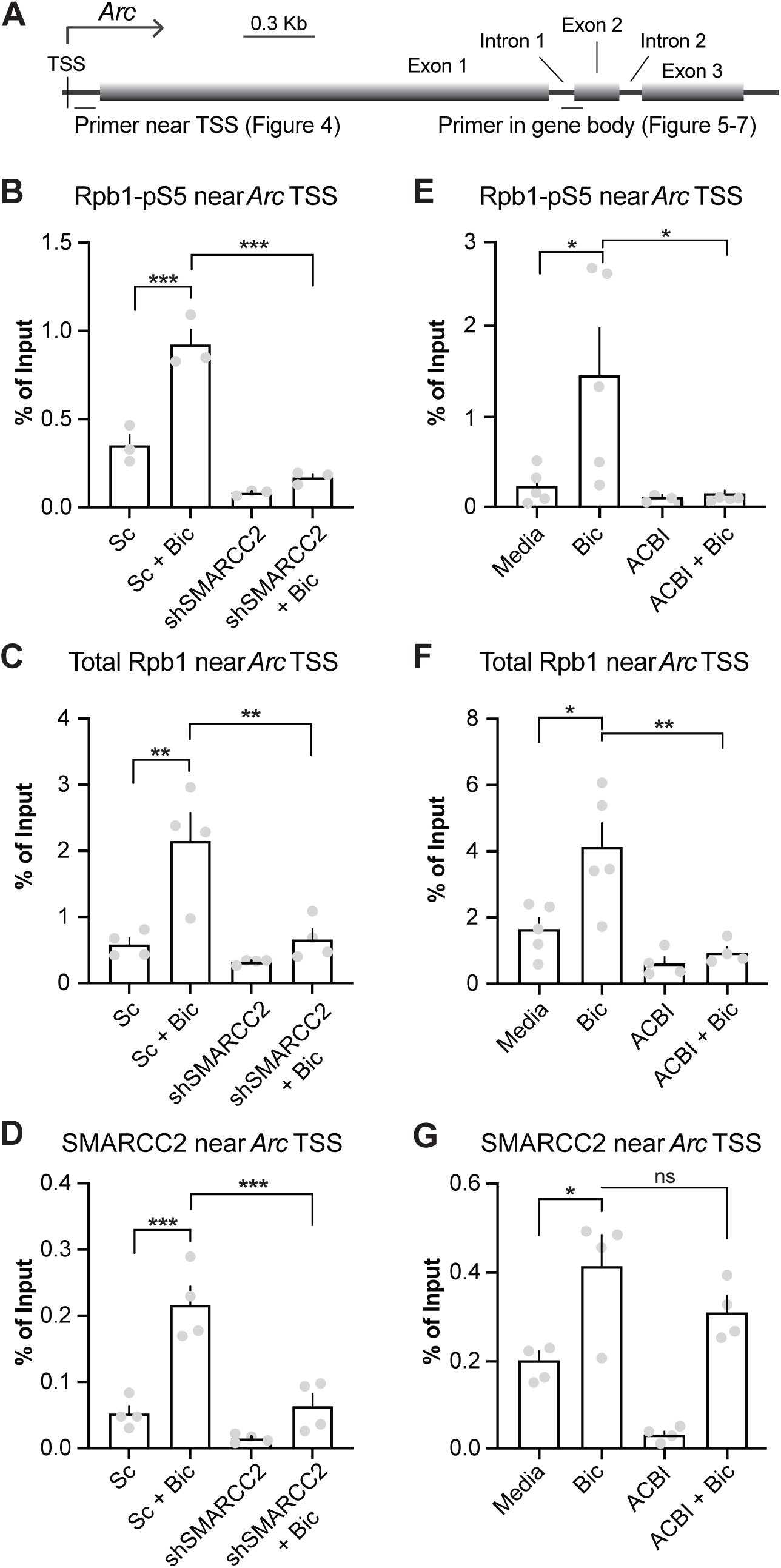
The nBAF complex aids activity-induced Pol2 recruitment to the *Arc* promoter. A) Schematic representation of *Arc* to show promoter, exons, introns, and primer positions used to quantified immunoprecipitated chromatin in this and other figures. TSS: transcription start site. B, C, and D) Neurons were SMARCC2 depleted by RNAi and knockdown was confirmed independent Western blots (not shown). Sc: Scrambled shRNA as control. E, F, and G) Neurons were incubated with ACBI (3 hours, 2.5 µM) to degrade SMARCA4. ChIP assays were subsequently performed for all treatments B-G. B and E) Quantified paused RNA Pol II binding near the *Arc* promoter determined by ChIP with an antibody against Rpb1 phosphorylated at Serine 5 in the CTD (Rpb1-pSer5). C and F) Quantified total RNA Pol2 binding near the *Arc* promoter determined by ChIP with antibody against Rpb1-NTD. D and G) Quantified SMARCC2 binding near the *Arc* promoter determined by ChIP with antibody against SMARCC2. Grey dots represent biological replicates; errors bars show SE of the mean. \**P*<0.05; \*\**P*<0.01; \*\*\**P*<0.001. One-way ANOVA analyses followed by Tukey’s *post hoc* tests were performed.

### The nBAF complex interacts with and is regulated by the Pol2 elongation complex (EC)

The purpose of activity-induced SMARCC2 translocation to the promoter region, we postulated, is to engage with the Pol2 EC and assemble the nBAF complex. A key EC component is P-TEFb, which signal-dependently switches Pol2 from a promoter proximal paused state to productive elongation by phosphorylating, among other targets, position two serine residues of Pol2 Rpb1 CTD heptads (Rpb1-pS2)^68^. To test if the BAF complex interacts with EC, we performed a Co-IP with CDK9, the catalytic subunit of p-TEFb. SMARCA4 and SMARCC2 was found to interact with CDK9, where the level of such interactions increased in response to activity (Figure 5A). As a control, for similar treatments, there were no changes in interactions of CDK9 with SPT6, another key member of the Pol2 EC^69^. Interestingly, SMARCC2 interacted with CDK9 despite ACBI1-mediated degradation of BAF subunits such as SMARCA4 (Figure 5A). This observation echoes previously stated activity-induced independent translocation of SMARCC2 to the *Arc* promoter region (Figure 4G) and suggests that CDK9-SMARCC2 may serve as the seed complex for activity-induced nBAF accretion. To test this possibility, we depleted CDK9 using the recently described monomeric CDK9 PROTAC (Thal-SNS032)^70^ and studied activity-induced nBAF assembly. Thal-SNS032 treatment for three hours depleted CDK9 levels (data not shown). CDK9 depletion attenuated activity-induced enhanced interaction of SMARCC2 with ARID1A, but not SS18L1 (Figure 5B), suggesting that partial nBAF assembly can occur without the kinase, but physical presence of CDK9 is necessary for activity-triggered complete assembly of nBAF. We also curiously noted that CDK9 degradation increased interactions between SMARCC2 and SS18L1 in unstimulated neurons, suggesting a nuanced role of the kinase in the process.

**Figure 5.**
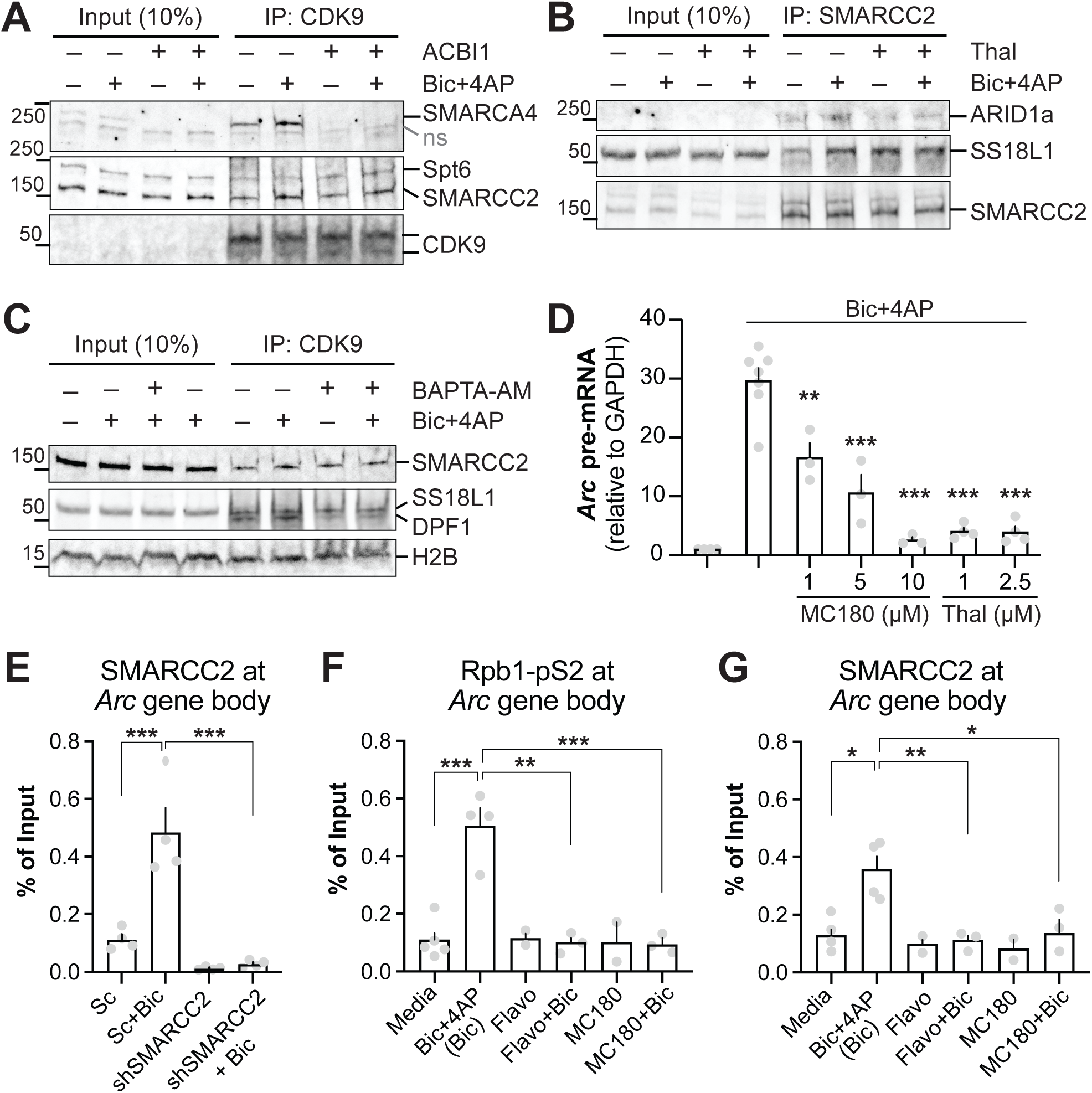
The nBAF complex interacts with and is regulated by Pol2 EC. A, B, and C) Co-IP was performed using anti-CDK9 (A and C), anti-SMARCC2 (B). Neurons were incubated for three hours with 2.5 µM with ACBI1 (A) or Thal-SNS-032 (B) and activity was induced by Bic+4AP for 15 minutes. Co-IP was then performed with cell lysates. C) Neurons were pre-treated with BAPTA-AM (100 µM, 15 minutes), then treated with Bic+4Ap for another 15 minutes, and lysates were co-immunoprecipitated. Co-IP material was electrophoresed and Western blotted to detect interacting subunits. Approximate position of nearest molecular weight marker is depicted. D) Neurons were incubated with MC180 and Thal-SNS-032 for 20 minutes and three hours respectively, followed by activity induction for 15 minutes with Bic+4AP. Normalized *Arc* pre-mRNA levels were assayed and is displayed. E) Quantified elongating RNA Pol II binding inside the *Arc* gene body, after 15 minutes of Bic+4AP stimulation, determined by ChIP with antibody against Rpb1-pSer2. F and G) Quantified SMARCC2 binding inside the *Arc* gene body, after identical stimulation as E, determined by ChIP in SMARCC2 depleted cells or in neurons treatments with MC180 or Flavopiridol (Flavo). \**P*<0.05; \*\**P*<0.01; \*\*\**P*<0.001. One-way ANOVA analyses were performed followed by Tukey’s *post hoc* test.

How does CDK9 facilitate activity-induced nBAF assembly? CDK9 is known to be regulated by Ca^2+^ signaling^71^; Ca^2+^ signaling is also necessary for nBAF assembly (Figure 3D). We wondered, if these two events are connected. Preventing Ca^2+^ signaling via BAPTA-AM impaired activity-induced association of CDK9 with the nBAF subunit DPF1 (Figure 5C). Together, our data suggests that synaptic activity induces nBAF assembly via Ca^2+^ signaling and requires CDK9.

Next, we inquired the effect of CDK9–nBAF association on *Arc* transcription. Induced *Arc* transcription, as expected^51,72^, was dose-dependently and highly sensitive to MC180^73^ (catalytic inhibition) or degradation of CDK9 (Figure 5D). Next, given that the EC travels with Pol2 in gene body during productive elongation^74^ and activity induces nBAF-CDK9 association, we asked if the BAF complex also traverses the gene body during ongoing transcription. ChIP with anti-SMARCC2 revealed that compared to control, activity induced enrichment of the subunit in *Arc* gene body (Figure 5E). Attesting to signal specificity, such enrichment was not seen in neurons depleted of the subunit. Next, we treated neurons with CDK9 inhibitors. As expected, elongation competent Rpb1-pS2 levels increased in *Arc* gene body in response to activity but dropped to baseline in response to CDK9 inhibitors (Figure 5F). Similarly, activity-induced enrichment of SMARCC2 in the gene body was also sensitive to CDK9 inhibition (Figure 5G). Taken together, these datasets suggest that SMARCC2 likely accompanies the EC through the gene body during productive elongation.

### The nBAF complex interacts with elongation-competent Pol2

If the BAF complex is to have a regulatory role in transcription elongation, its signal-dependent interaction with elongation-competent Pol2 is highly likely. We tested this possibility by performing Co-IP with an antibody against Rpb1-pS2. The interactions between Rpb1-pS2 increased with SMARCC2 and SS18L1 in response to activity (Figure 6A). This enhanced interaction was sensitive to inhibitors of CDK9 activity, which globally block productive elongation^72^ (Figure 6A). Next, we used BD98 to prevent nBAF assembly (Figure 2F) and studied interactions of Rpb1-pS2 with various BAF subunits. BD98 hindered interaction of Rpb1-pS2 with nBAF subunits such as ARID1A, as well as SMARCA4 and SMARCC2 (Figure 6B). To test if the BAF complex could be detected within the gene body during active transcription, we performed ChIP. Activity significantly enriched SMARCC2 levels in the *Arc* gene body (Figure 6C), which aligns with our Co-IP data above that demonstrate interactions of the nBAF complex with EC and elongation-competent Pol2. Such activity-induced SMARCC2 gene body enrichment was sensitive to the nBAF inhibitor BD98 and BAF degrader ACBI1 (Figure 6C). Notably, activity-induced SMARCC2 enrichment was differentially susceptive of ACBI1 at the *Arc* promoter (insensitive; figure 4G) versus in the gene body (sensitive). Together, data presented so far suggest a promoter proximal modular assembly of nBAF, followed by its participation in Pol2 productive elongation.

**Figure 6.**
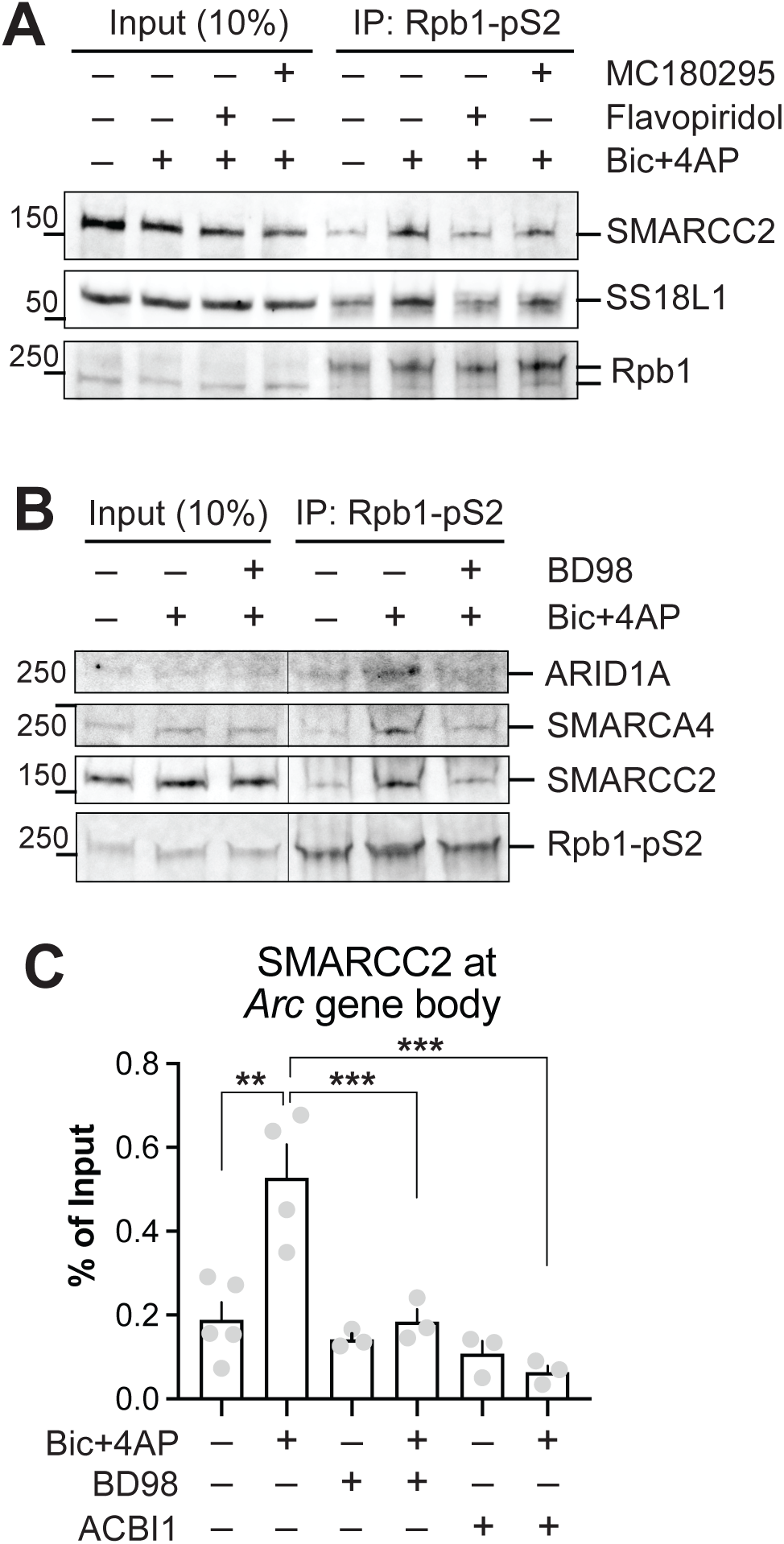
The nBAF complex interacts with elongation competent Pol2. A) Neurons were incubated with CDK9 inhibitors (MC180295 and Flavopiridol, 5 µM) for 20 minutes and neuronal activity was then induced with Bic+4AP for 15 minutes. Co-IP was performed with Rpb1-pS2 antibody that recognizes elongation competent Pol2. Interactions between elongating Pol2 and BAF subunits SMARCC2 and SS18L1 were tested by Western blotting the immunoprecipitates. Total Rpb1 is displayed as loading control. B) Neurons were treated as in A, except, they were incubated with BD98 (20 minutes, 5 µM). Interactions between Rpb1-pS2 and indicated nBAF subunits were verified. C) ChIP data demonstrating SMARCC2 binding inside the *Arc* gene body in response to 15 minutes of activity in the presence or absence of BD98 or ACBI1. Grey dots represent biological replicates; errors bars show SE of the mean. \**P* <0.05; \*\**P*<0.01; \*\**P*<0.001. One-way ANOVA analyses were performed followed by Tukey’s *post hoc* test.

### Assembled complex and its ATPase activity is required for productive elongation

To decouple the role of nBAF in productive elongation from its involvement in Pol2 recruitment and initiation at the promoter, we used Triptolide. Triptolide is an inhibitor of the ATPase activity of XPB, the TFIIH helicase/translocase subunit found in the transcription initiation complex. Triptolide treatment prevents new transcription initiation but doesn’t interfere with promoter escape and productive elongation of paused Pol2^37,75,76^. To verify its effect on *Arc* transcription, we treated neurons with Triptolide and subjected them to activity. Transcription was significantly induced in Triptolide-treated neurons, but as expected due to obstructed Pol2 initiation, was less compared to the full-bodied transcription in counterpart neurons without the inhibitor (Figure 7A). Strong induction despite XPB/TFIIH inhibition indicates the ability of promoter proximal paused Pol2 to enter productive elongation signal-dependently. Such initiation-independent elongation was next studied after inhibiting induced nBAF assembly (BD98) or its ATPase activity (BRM014). Both BD98 and BRM014 significantly reduced initiation-decoupled *Arc* transcription (Figure 7A). These data strongly suggest that nBAF complex is necessary for productive elongation of *Arc*.

**Figure 7.**
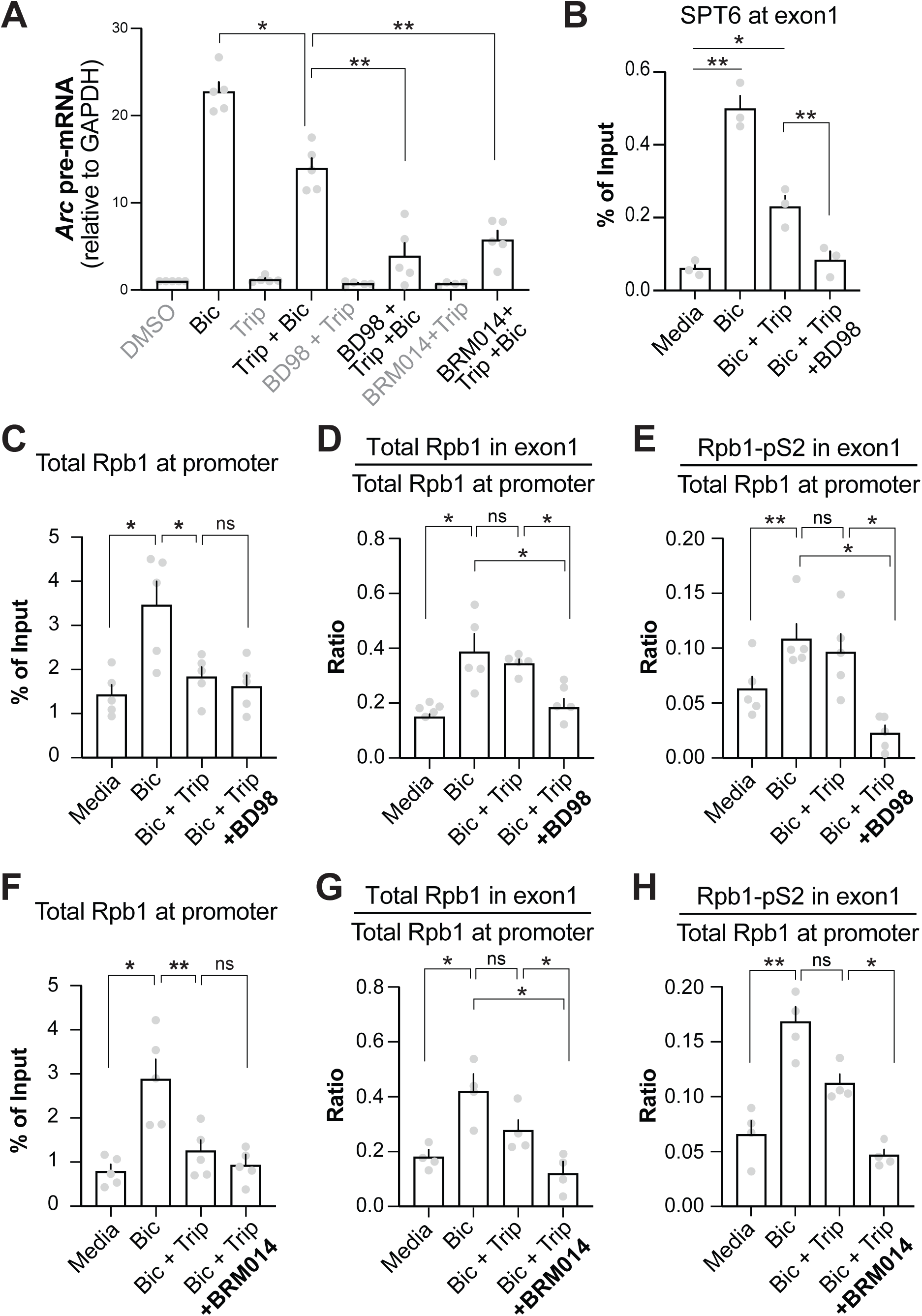
Assembled complex and its ATPase activity is required for Pol2 productive elongation. A) Neurons, as indicated, were incubated with BD98 (5µM) or BRM014 (5µM) for 20 and 10 minutes respectively and cotreated with Triptolide (1µM) and Bic+4AP for 10 minutes. Activity-induced *Arc* transcription was quantified by *Arc* pre-mRNA levels (normalized by *GAPDH* pre-mRNA). B-E) Neurons were incubated with BD98 for 20 minutes and cotreated with Triptolide and Bic+4AP as indicated. ChIP assays were performed with antibodies against SPT6, Rpb1-NTD, and Rpb1-pS2. Levels of SPT6 in *Arc* gene body is displayed in (B). Total Pol2 levels at *Arc* promoter is displayed in (C). Total Pol2 levels in the gene body, normalized by total Pol2 at the promoter, is shown in (D). Rpb1-pS2 levels in gene body, normalized by total Pol2 at the promoter, is displayed in (E). F-H) In a parallel set of experiments, neurons were treated as in B-E, except they were incubated with BRM014 prior to experiencing activity. Total Pol2 levels at *Arc* promoter is displayed in (F). Total Pol2 levels in the gene body, normalized by total Pol2 at the promoter, is shown in (G). Rpb1-pS2 levels in gene body, normalized by total Pol2 at the promoter, is displayed in (H). Grey dots represent biological replicates; errors bars show S1E of the mean. \**P*<0.05; \*\**P*<0.01; \*\**P*<0.001. One-way ANOVA statistical analyses were performed followed by Tukey’s *post hoc* test.

To corroborate the relationship of nBAF and Pol2 elongation further, we performed a series of ChIP assays focusing on the *Arc* gene body. First, we assayed for SPT6, a representative of the EC. SPT6 levels significantly increased in response to activity, both in the absence and presence of Triptolide, albeit less so in the latter (Figure 7B). Echoing gene transcription trends seen in figure 7A, such activity-induced SPT6 gene body enrichment was attenuated when neurons were also treated with BD98 (Figure 7B). Next, two sets of ChIP assays were performed with and without Triptolide using antibodies against total Rpb1 and elongation competent Rpb1-pS2. Total Rpb1 near the promoter was used to normalize amounts of Rpb1 and Rpb1-pS2 in the gene body. This normalization was necessary because activity-induced promoter proximal Rpb1 levels differed in the presence of Triptolide (Figure 7C and 7F). When normalized as above, levels of Rpb1 and Rpb1-pS2 in the gene body were comparable in neurons with and without Triptolide. Also, their levels were significantly more after activity versus control (Figure 7D, 7E, 7G, and 7H). To unveil underlying mechanisms, neurons were also treated with BD98 to block nBAF assembly and BRM014 to inhibit ATPase activity during initiation-decoupled elongation. Both small molecule interventions attenuated activity-induced enrichment of Rpb1 and Rpb1-pS2 in *Arc* gene body (Figure 7D, 7E, 7G, and 7H), demonstrating that productive elongation of Pol2 in *Arc* gene body requires activity-induced assembly of nBAF and its ATPase activity.

Finally, to broaden the scope of nBAF function in rIEG transcriptional elongation and to rule out above observations as only an *Arc*-specific phenomenon, we extended the ChIP assay to five additional rIEGs (*cFos, Gadd45g, Cyr61, Btg2,* and *Dusp1*), which displayed sensitivity to ACBI and BD98 (Figure S1). The trend of Rpb1-pS2 levels in the body of these genes was like that seen above in *Arc*; levels increased after Bic4AP treatment and activity, remained comparable (or, decreased somewhat) if elongation was isolated from initiation, but dropped significantly if nBAF assembly was inhibited with BD98 along with initiation-elongation decoupling (Figure S3). Taken together, our data demonstrate that nBAF regulates productive elongation in rIEG gene bodies via interaction with the elongation complex and elongation-competent Pol2, which is mechanistically mediated by activity-induced assembly and its ATPase activity.

## Discussion

This study contributes the following novel observations: 1) neuronal activity induces nBAF assembly, 2) activity-assembled nBAF mediates Pol2 productive elongation, and therefore, 3) nBAF is necessary for activity-induced rIEG transcription.

The BAF complex is often thought of as a developmentally assembled polymorphic complex, which undergoes a cell-type specific compositional transformation in neurons, referred to as the nBAF^18^. It remained unknown if nBAF is a steady-state complex that remains compositionally impervious to neuronal activity, or if it can undergo recurring rounds of complex formation, especially in response to transcriptional profile-altering cellular signals. Here, we show that neuronal activity induces ancillary assembly of nBAF on top of its basal level (Figure 2F). Our data indicate, such assembly is ordered and modular in nature. Modular assembly of BAF complex was recently described^14^, where complex formation is initiated by the SMARCC1/SMARCC2 core module. This core binds next to the subunit defining ARID1 module, which then binds to the BRG1/BRM and SS18/L1 containing ATPase module to finalize the assembly. The following observations led us to believe that activity-induced neuronal nBAF assembly is similarly modular: 1) inhibition of ARID1A– SMARCC2 binding with BD98 also attenuates SMARCA4–SMARCC2 association (Figure 2F), 2) SS18L1 assembles with SMARCC2 only in the presence of the ATPase (Figure 3E), and 3) SMARCC2 is independently recruited to *Arc* promoter region and binds to EC (CDK9) despite ACBI1-dependent depletion of SMARCA4 (Figure 4G and 5A). Furthermore, during the preparation of this manuscript, an independent study in preprint^77^ demonstrated activity-induced enhanced interaction of SMARCA4 with ARID2 (activity-assembled PBAF?). Considered together with our study, it is possible that membrane depolarization induces assembly of both nBAF and PBAF, which then likely undertake independent and overlapping functions to mediate the span of nuclear responses to activity.

The relationship between the BAF complex and RNA Pol2 has been studied over the years in many models and cell types. These studies include genome-wide studies where ChIP with high-throughput sequencing (ChIP-seq) and other global approaches suggest that BAF subunits largely enrich at gene promoters and enhancers, where they facilitate functions of these genomic regions^33,37,78–83^. From these studies, the BAF complex may be inferred to have promoter- and enhancer-specific functions only. Pol2 elongation, being dynamic and non-synchronous in a cell population, can be elusive to capture with global techniques such as ChIP-seq of BAF subunits. Therefore, we approached the issue using rIEGs at an early timepoint where paused Pol2 release, rather than recruitment, primarily mediate transcriptional changes. Such instances lend themselves well to decoupling of productive elongation from Pol2 recruitment and initiation via small molecule pharmacology (e.g., Triptolide). Our Triptolide assays with *Arc* and other rIEGs (Figure 7 and S3) clearly show the necessity of nBAF during productive elongation. During elongation, the nBAF complex works in conjunction with the elongation complex, perhaps boosting each other’s functions. Whether nBAF is also responsible for Pol2 elongation in other longer genes remains to be investigated in the future with genome-scale approaches such as mammalian NET-seq^84^, a powerful technique that still needs optimization in non-dividing cells such as neurons.

In lieu of its role in Pol2 elongation, nBAF is necessary for activity-induced transcription of rIEGs. This inference is drawn from our above studies that utilized RNAi against two BAF subunits, two degraders of various BAF complexes, and inhibitors of nBAF assembly or its catalytic activity. However, our findings appear to contradict a previous study^85^, which reported that loss of neuron-specific BAF subunit ACTL6B derepresses rIEGs. There are several avenues to reconcile with apparent differences in our conclusions. One, compared to our inhibitors and degraders that work in minutes to hours, RNAi and knocking out a target in animals — as used by the previous study — deplete the protein of interest over days. Such longer durations leave room for cellular homeostatic mechanisms (e.g., retention of the non-neuronal paralogue ACTL6A instead^85^) to influence the outcome. Such possibilities are supported by our observation in figure 1 where ACBI1 treatment for three hours degrades ACTL6B but does not derepress rIEG transcription. Two, ACTL6A is a constituent of cBAF, PBAF, and GBAF. It is possible that ACTL6B, the neuronal paralogue, similarly convenes with all three BAF complexes in neurons and the reported phenotypic outcome of its loss is underwritten by one or more malfunctioning BAF complexes. Taken together, the role of BAF complex in activity-induced rIEG is nuanced, where complex subtype and its composition likely play complementary or contradictory roles in a context-dependent fashion. To comprehend such nuanced set of functions elaborately, future work must be directed to distinct BAF subcomplexes.

Our current findings will be relevant to several human disorders and diseases that, on one hand, are associated with inefficient gene transcription, especially aberrant productive elongation, and on the other hand, correlate with mutations in genes encoding BAF subunits. For example, among non-neuronal cells, many forms of cancer stem from defective Pol2 elongation^86–88^. Many of these cancers also prominently feature mutations in BAF genes^89^. In the brain, anomalies of Pol2 elongation have been implicated in NDDs^90^ and brain cancers^91^. With regards to NDDs – ‘BAFopathies’ – genes coding for cBAF subunits possess most *de novo* missense and protein-truncating mutations among all nuclear protein complexes^28^. Also, biallelic mutations in the neuron-specific *ACTL6B* or haplosufficiency of *ARID1B* causes recessively inherited autism^24,85^, microduplication of *ARID1A* causes intellectual disability^92^, and *SMARCC2* is a high-confidence autism gene^93–97^. As for brain tumors, *SMARCA4* is recurrently mutated in multiple cancers^98^. Future studies are expected to reveal cause-effect association between these mutations, Pol2 elongation defects, and human maladies.

In summary, we have demonstrated that nBAF is signal-dependently assembled, which then allows the complex to mediate several aspects of induced Pol2 activity at gene promoters and gene bodies. Such assembly likely occurs at or near promoters and requires interactions with pTEFb. Association with pTEFb likely is part of a larger web of interactions that include elongation competent Pol2. It remains to be tested if the nBAF complex is a steady state constituent of the Pol2 elongation complex or an intermittent participant that aids the elongation process to overcome nucleosomal barriers and/or undertake co-transcriptional splicing as needed. It also remains to be investigated if PBAF and GBAF undertake synergistic roles in activity-induced IEG transcription or of other longer genes implicated in human disorders. Among these longer genes, it will be of interest to investigate if BAF complexes have unique or redundant roles in regulating Pol2 velocity and it passage through chromatin barriers. Last but not the least, we look forward to further development of BAF complex inhibitors and degraders to make them blood brain barrier permeable, which will then allow extension of our current enquiries to the brain.

## Acknowledgments

Technical support by Baani Minhas, Courtny Nimez, and Isabel Ramos is acknowledged. This study was funded by the following National Institutes of Health grants: 1) from the National Institute of Environmental Health Sciences (NIEHS) to RNS (R01ES028738), and 2) from the National Institute of Mental Health (NIMH) to RNS and ECD (R21MH128678).

## Author contributions

KGC, AV, and MHS, performed experiments, curated, and validated data, performed formal analyses, and edited the manuscript. KGC co-wrote the first draft. MD, BGR, LBK, SGN performed experiments and acquired data. ECD provided reagents, expertise, and feedback. RNS conceptualized, supervised, and administered the project, performed experiments, visualized figures, and wrote the first draft. RNS and ECD acquired funding. All authors reviewed the manuscript.

## Methods

### Primary neuronal culture

Experiments were performed using dissociated cortical rat neurons obtained from Sprague-Dawley rats. Time-pregnant rats were purchased from Charles River Laboratories. Rats were delivered (E18) 2 days prior to dissection and housed individually on a 12-hour light/dark cycle with access to food and water *ad libitum*. Dams were anesthetized with Euthasol^®^ (390 mg/ml sodium pentobarbital and 50 mg/ml sodium phenytoin) and decapitated using a guillotine. Embryonic day 18 pups were removed and used to prepare primary cultures of cortical neurons under animal protocol #22-1143, reviewed and approved by the University of California, Merced IACUC. To remove cortical hemispheres, heads were cut and transferred to a plate containing HBSS plus Ca^2+^and Mg^2+^ (Gibco, catalog no. 14025092). A medial cut was made caudal to rostral of the head and gently pushed the brain out without disturbing the cortex. Under a dissecting microscope, the brain was placed with the ventral surface facing upwards, a cut was made in the sagittal plane to separate the hemispheres. Meninges were then carefully removed thoroughly. The collected tissue was placed in a dish containing HBSS plus Ca^2+^and Mg^2+^ (Gibco, catalog no. 14025092) and taken to a tissue culture room for mechanical dissociation. 1 milliliter of StemPro^®^ Accutase^®^ (Life Technologies, Inc., catalog no. A1110501) was added for a 7-minute digestion, followed by mechanical dissociation with fire-polished Pasteur pipettes. The reaction was stopped by adding 5 milliliters of HBSS lacking Ca^2+^and Mg^2+^ (Gibco, catalog no. 14175095). The dissociated neuron resuspension was then centrifuged for four minutes at 200 rcf, resuspended in plating media, and used to do a cell count.

Cell counts were done using trypan blue and a TC20TM automated cell counter (Bio-Rad, catalog no. 1450102), and subsequently plated to dishes containing pre-warmed Neurobasal medium (Gibco, catalog no. 21103049) supplemented with 25 µM glutamate (Sigma-Aldrich, catalog no. 1446600), 0.5 mM L-glutamine (Sigma-Aldrich, catalog no. G8540). 100 cm^2^ dishes were plated at a density of 7.8 x 10^6^ neurons or 35 cm^2^ dishes at a density of 1.0 x 10^6^. Cells were maintained at 37°C in a humidified incubator with 5% CO2. Neurons were grown for up to two weeks in the medium described above without glutamate, replacing half the media every 3– 4 days. Matured neurons were then used for various assays between 10-14 days..

### Pharmacological Treatments

Neuronal activity was induced by co-treating neurons with 50uM Bicuculline (Sigma-Aldrich, catalog no. 14340) and 75 µM 4-aminopyridine (Acros Organics, catalog no. 104571000). To degrade BAF complex, neurons were treated with ACBI1 (OpnMe). 1µM or 2.5µM ACBI1 were used for Western blot analysis and RNA assays. 2.5 µM ACBI1 dose was used for ChIP assays. Cells were pre-treated for 3 hours. SMARCA4 protein degradation was used as a control to assess ACBI1 efficiency. cisACBI1 (OpnMe) was used as negative control. To inhibit BAF ATPase function, BRM014 (compound 14, MedChemExpress; catalog no. HY-119374) and FHT-1015 (MedChemExpress; catalogue no. HY-144896) were used at concentrations described in figure legends. BD98 (BRD-K98645985), BRD7i 2-77 (PBAF inhibitor), BRD9i (GBAF inhibitor), and VZ-185 (PBAF and GBAF degrader) were obtained from the Dykhuizen lab. BD98 is now commercially available from MedChemExpress (catalogue no. HY-114268). BRD7i, BRD9i, and VZ-185 are commercially available from OpnMe. Cells treated with BRM014, FHT-1015, BD98, BRD7i 2-77, or BRD9i were pre-treated for 15-30 mins at concentrations indicated in figures, with or without Bicuculline+4AP added in the last 15 minutes of the treatment. VZ-185 (0.25 µM) was pre-treated for three hours with or without Bicuculline+4AP added in the last 15 minutes. Brd7 and/or Brd9 protein degradation was used as a control to assess VZ-185 efficiency. To inhibit phosphorylation of RNA Pol II CTD (pS2), MC180295 and Flavopiridol were used at indicated concentrations. Cells were treated for 20 minutes with the inhibitors, which was followed by 15 minutes of induced neuronal activity with Bicuculline+4AP. To degrade CDK9, PROTAC Thal-SNS032 was used. Cells were pre-treated for 3 hours with or without induced neuronal activity with Bicuculline/4AP. RNA assays utilized 1µM or 2.5 µM final concentrations. Co-IP cell treatments used 2.5 µM final concentrations. CDK9 protein degradation was used as a control to assess Thal-SNS032 efficiency. To inhibit neuronal firing but not synaptic activity, Tetrodotoxin was pre-treated at a concentration of 1 µM for five minutes. To inhibit glutamate-gated synaptic transmission, cells were treated with a cocktail composed of RS-CPP (2.5 µM), CNQX (2.5 µM) and Nimodipine (5 µM) for five minutes. To chelate intracellular Ca^2+^, neurons were pre-treated with BAPTA-AM (100 µM) for 15 minutes followed by induced neuronal activity with Bicuculline+4AP.

### Co-Immunoprecipitation (Co-IP)

After performing treatments as indicated, neurons were lysed in 335uL of IP buffer (5-10% Ficoll; 1mL glycerol; 150mM NaCl; 20 mM Tris–HCl [pH 7.5]; 1.25 mM EGTA; 2 mM EDTA; .05% Tween-20). Protease/phosphatase inhibitor cocktail was added as recommended. Cells were lysed 8-12 times with a 26-gauge needle and sheared by sonication (8 × 30s, HIGH setting in Bioruptor^®^). Sonicated material was centrifuged at 15,000 rpm for 1 min at 4 °C to clear debris. Antibodies (2-5µg) were pre-loaded on A/G beads for 30 minutes at 35°C and 5-10% inputs were set aside for each treatment. Sonicated material and preloaded magnetic A/G beads (Pierce) were pulled down for 2 hours at 35°C with constant rotation. A/G bead-protein complexes were prepared for gel electrophoresis by resuspending in sample buffer (4X Laemmli sample buffer with 10% ý-mercaptoethanol; diluted to 1X with IP buffer) and heating at 95°C for 5-10 minutes. Beads were separated by a magnetic rack and processed samples were electrophoresed. Co-IP assays were performed in 2-3 biological replicates.

### Gel Electrophoresis and Western Blotting

Neurons were lifted from cell culture dishes with sonication buffer (20 mM Tris–HCl [pH 8]; 1.25 mM EGTA; 2 mM EDTA; 0.5% SDS) supplemented with 1:100 protease/phosphatase inhibitor cocktail (Cell Signaling; catalog no. 5872S). Lysates were sheared by sonication (8 × 30s, HIGH setting in Bioruptor^®^). Sonicated material was centrifuged at 15,000 rpm for 1 min at 4 °C to clear debris.

Samples were combined with (4X Laemmli sample buffer; Bio-Rad; catalog no. 1610747) with 10% β-mercaptoethanol (Sigma; catalog no. 63689) and were warmed for 10 min at 95°C in a heat block. Sample and dye mixtures were then loaded in a 4-20% (Bio-Rad; catalog no.: 4568095) or 4-15% (Bio-Rad; catalog no.: 456-1083) gels. After electrophoresis at 90-100V, gels were transferred to a polyvinylidene difluoride (PVDF) membrane (Bio-Rad; catalog no. 10026933) using the Bio-Rad Trans-Blot Turbo Transfer System with 20% Methanol-containing transblot turbo transfer buffer (Bio-Rad; catalog no. 10026938) for 10-15 minutes. PVDF membranes were immediately transferred to cold Tris-buffered saline with Tween-20 (TBS-T 0.1%) and incubated at 4°C overnight in primary antibody solution [0.1% TBS-T supplemented with 3% BSA (Fisher; catalog no.: BP9703) and protease-phosphatase inhibitors]. Membranes were washed three times every 5 minutes in 0.1% TBS-T before being probed with secondary antibody for 60 minutes at room temperature.

Secondary antibodies were either goat-anti–mouse 647 (RRID: A21236) or goat-anti–rabbit 488 (RRID: A11034) Alexa Fluor secondary antibodies (Life Technologies). Membranes were washed three times with 0.1% TBS-T for 5 min each and imaged using Bio-Rad Multiplex ChemiDoc Imaging System. Densitometric quantifications were performed in Image Lab Software 2020.

### Chromatin Immunoprecipitation (ChIP)

Neurons were cross-linked with 1% formaldehyde for 5 minutes after indicated treatments. Crosslinking reaction was quenched for 5 minutes with 1.25 M glycine buffer in PBS. Cross-linked cells were resuspended in sonication buffer (20 mM Tris–HCl [pH 7.5]; 1.25 mM EGTA; 2 mM EDTA; 0.5% SDS, supplemented with inhibitors for proteases and phosphatases, 1:100) followed by sonication for 16 cycles (bursts and intervals of 30s each) with the Bioruptor^®^ (Diagenode), which produced 200-1000 bp genomic DNA fragments. Sonicated samples were then immunoprecipitated overnight at 4°C in IP buffer with 2-5μg antibody. 20% of the sonicated material was set aside for inputs. Antigen-antibody complexes were immunoprecipitated with Pierce Protein A/G Magnetic Beads, washed four times with low salt buffer, once with LiCl buffer, and finally once with Tris-EDTA. Samples were reverse cross-linked at 65°C degrees overnight, and chromatin DNA was eluted using the Thermo Scientific™ GeneJET Plasmid Maxiprep Kit. Eluted chromatin was quantified by qPCR. Heat map in figure S3 was generated using ChIP material used figure 7C-E. Primers against promoter and gene body (∼1000 bp downstream of TSS) of five rIEGS were used to create the heat map. In summary, total Rpb1 near the promoter was used to normalize amounts Rpb1-pS2 in the gene body for each treatment. Heat map depicts the mean value of *N*=3 with a baseline-correction (baseline is the Media treatment) calculated as a percentage (100 * (value/baseline).

### RNA-based pre-mRNA quantification assays

Total RNA was extracted using the RNAspin Mini kit (GE Lifesciences; catalog no. 25050072) following the manufacturer’s instructions. RNA processing, PCR amplification and quantification has been previously described in details^99^. The Bio-Rad CFX Connect real-time PCR Detection System was used here.

### Cellular Thermal Shift Assay (CETSA)

CETSA was adapted from Jafari *et. al.*^60^. Briefly, neurons were treated with BD98 (20 μM) or DMSO as a control. Cells were scraped, collected in 0.5ml of media, and centrifuged at 500 rcf. Supernatant was removed and cells were resuspended in 0.5ml of 1x PBS. For each treatment, cell lysate was split and a portion was added to PCR tubes. Each tube was individually subjected to a unique temperature. We initially tried a temperature range of 37°C – 47°C and found BAF subunits degrading sufficiently at 43.3°C. Samples were exposed to 37°C or 43.3°C temperate for three minutes, then left at room temperature for an additional three minutes. Then samples were frozen for 12 minutes at -80 degrees, fast thawed, and frozen again. After freeze/thawing, samples were placed on ice and transferred to clean. Transferred lysates were centrifuged at 15000 rpm for 20 minutes at 4°C to remove debris and the supernatant was prepared for Western blot analysis.

### RNAi

SMARCC2 shRNA is previously published^58^. The shRNA construct for *ARID1A* (target sequence: TGGACCTCTATCGCCTCTATG, NM_006015) was obtained from Sigma (TRCN0000358749). This pLKO.1-based construct was packaged into lentiviruses. Self-inactivating HIV lentivirus particles were produced by transfecting 293 T cells with the shRNA vector, envelope (pMD2.G; Addgene), and packaging plasmids (psPAX2; Addgene) using a previously described protocol^99^. Efficiency of this construct was validated by Western blotting for ARID1A.

### Quantification and Statistical Analyses

Statistical analyses were conducted using GraphPad Prism 10 (GraphPad software, San Diego, CA). Error bars represent standard error of the mean. No tests for outliers were conducted; therefore, all data points were included. Effects were determined by *t* test or one-way ANOVA with appropriate *post hoc* tests for determination of *P* values; details are indicated in the figure legends. Biological replicates are indicated throughout within figures or in their legends. Biological replicates constitute cell culture preparations from the pooled cortices of embryos from independent dams.

### Antibodies

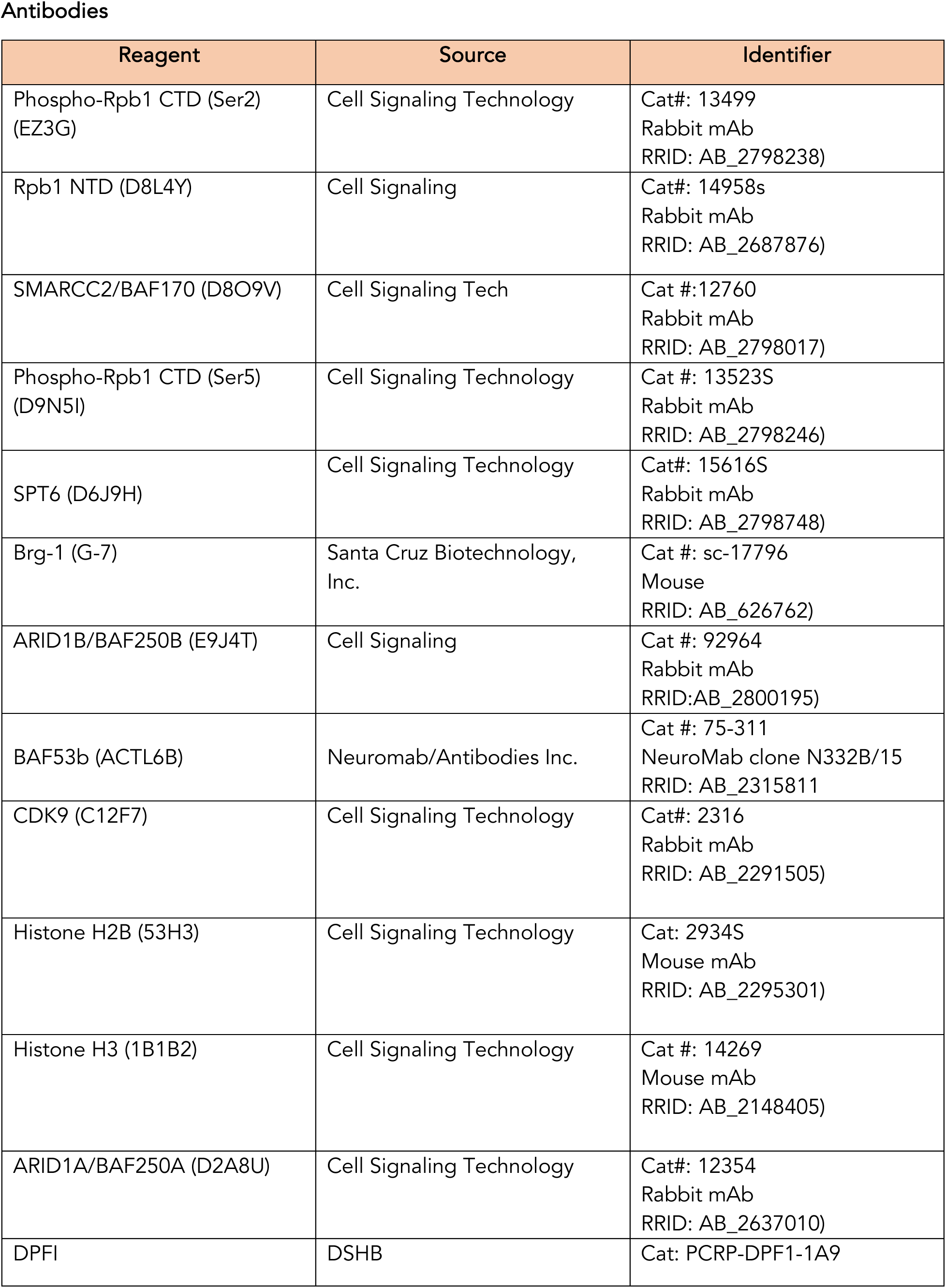
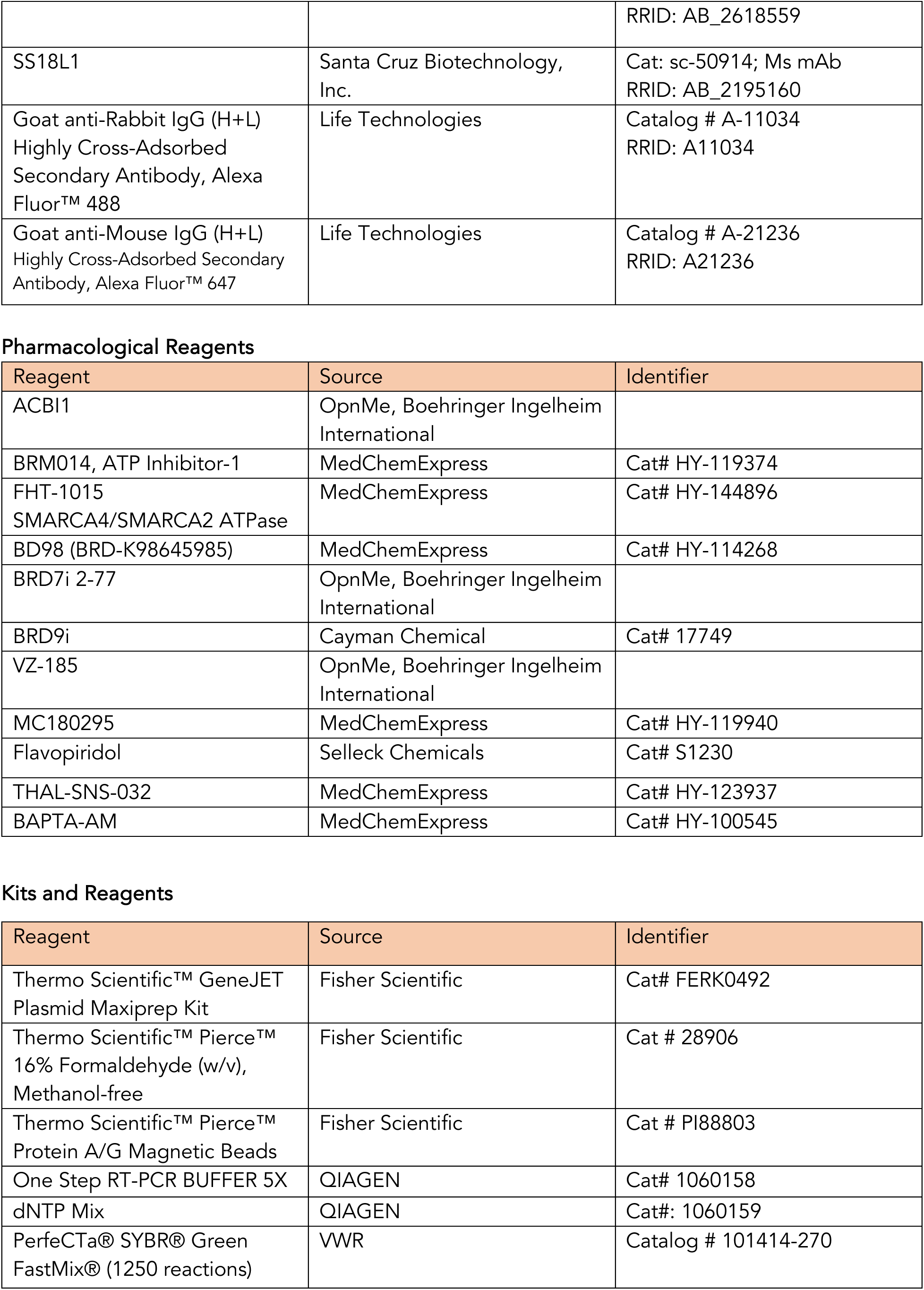
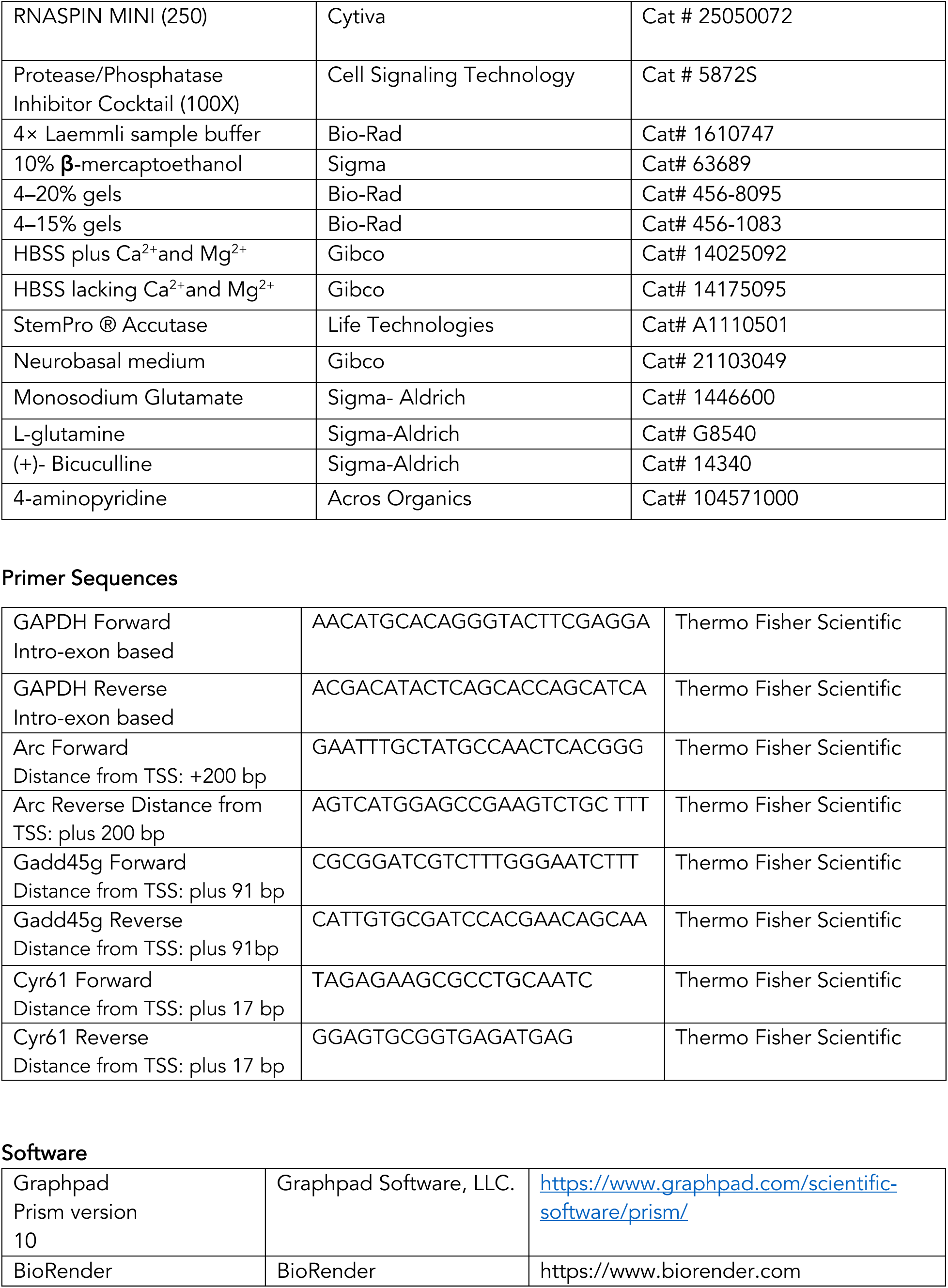

## References

1. Stern, M., Jensen, R., and Herskowitz, I. (1984). Five SWI genes are required for expression of the HO gene in yeast. J. Mol. Biol. 178, 853–868. 10.1016/0022-2836(84)90315-2.

2. Cairns, B.R., Kim, Y.J., Sayre, M.H., Laurent, B.C., and Kornberg, R.D. (1994). A multisubunit complex containing the SWI1/ADR6, SWI2/SNF2, SWI3, SNF5, and SNF6 gene products isolated from yeast. Proc. Natl. Acad. Sci. U. S. A. 91, 1950–1954. 10.1073/pnas.91.5.1950.

3. Neigeborn, L., and Carlson, M. (1984). GENES AFFECTING THE REGULATION OF SUC2 GENE EXPRESSION BY GLUCOSE REPRESSION IN SACCHAROMYCES CEREVISIAE. Genetics 108, 845–858. 10.1093/genetics/108.4.845.

4. Dingwall, A.K., Beek, S.J., McCallum, C.M., Tamkun, J.W., Kalpana, G. V, Goff, S.P., and Scott, M.P. (1995). The Drosophila snr1 and brm proteins are related to yeast SWI/SNF proteins and are components of a large protein complex. Mol. Biol. Cell 6, 777–791. 10.1091/mbc.6.7.777.

5. Wang, W., Côté, J., Xue, Y., Zhou, S., Khavari, P.A., Biggar, S.R., Muchardt, C., Kalpana, G. V, Goff, S.P., Yaniv, M., et al. (1996). Purification and biochemical heterogeneity of the mammalian SWI-SNF complex. EMBO J. 15, 5370–5382. 10.1002/j.1460-2075.1996.tb00921.x.

6. Wang, W., Xue, Y., Zhou, S., Kuo, A., Cairns, B.R., and Crabtree, G.R. (1996). Diversity and specialization of mammalian SWI/SNF complexes. Genes Dev. 10, 2117–2130. 10.1101/gad.10.17.2117.

7. Alpsoy, A., and Dykhuizen, E.C. (2018). Glioma tumor suppressor candidate region gene 1 (GLTSCR1) and its paralog GLTSCR1-like form SWI/SNF chromatin remodeling subcomplexes. J. Biol. Chem. 10.1074/jbc.RA117.001065.

8. Centore, R.C., Sandoval, G.J., Soares, L.M.M., Kadoch, C., and Chan, H.M. (2020). Mammalian SWI/SNF Chromatin Remodeling Complexes: Emerging Mechanisms and Therapeutic Strategies. Trends Genet. 10.1016/j.tig.2020.07.011.

9. Phelan, M.L., Sif, S., Narlikar, G.J., and Kingston, R.E. (1999). Reconstitution of a core chromatin remodeling complex from SWI/SNF subunits. Mol. Cell 3, 247–253. S1097-2765(00)80315-9 [pii].

10. Lessard, J., Wu, J.I., Ranish, J.A., Wan, M., Winslow, M.M., Staahl, B.T., Wu, H., Aebersold, R., Graef, I.A., and Crabtree, G.R. (2007). An essential switch in subunit composition of a chromatin remodeling complex during neural development. Neuron 55, 201–215. S0896-6273(07)00452-7 [pii].

11. Nguyen, H., Sokpor, G., Pham, L., Rosenbusch, J., Stoykova, A., Staiger, J.F., and Tuoc, T. (2016). Epigenetic regulation by BAF (mSWI/SNF) chromatin remodeling complexes is indispensable for embryonic development. Cell Cycle 15, 1317–1324. 10.1080/15384101.2016.1160984 [doi].

12. Narayanan, R., Pirouz, M., Kerimoglu, C., Pham, L., Wagener, R.J., Kiszka, K.A., Rosenbusch, J., Seong, R.H., Kessel, M., Fischer, A., et al. (2015). Loss of BAF (mSWI/SNF) Complexes Causes Global Transcriptional and Chromatin State Changes in Forebrain Development. Cell Rep. 13, 1842–1854. 10.1016/j.celrep.2015.10.046 [doi].

13. Ronan, J.L., Wu, W., and Crabtree, G.R. (2013). From neural development to cognition: unexpected roles for chromatin. Nat. Rev. 14, 347–359. 10.1038/nrg3413 [doi].

14. Mashtalir, N., D’Avino, A.R., Michel, B.C., Luo, J., Pan, J., Otto, J.E., Zullow, H.J., McKenzie, Z.M., Kubiak, R.L., St. Pierre, R., et al. (2018). Modular Organization and Assembly of SWI/SNF Family Chromatin Remodeling Complexes. Cell 175, 1272–1288.e20. 10.1016/j.cell.2018.09.032.

15. Ho, L., and Crabtree, G.R. (2010). Chromatin remodelling during development. Nature 463, 474–484. 10.1038/nature08911.

16. Yoo, A.S., Staahl, B.T., Chen, L., and Crabtree, G.R. (2009). MicroRNA-mediated switching of chromatin-remodelling complexes in neural development. Nature 460, 642–646. 10.1038/nature08139 [doi].

17. Yoo, A.S., Sun, A.X., Li, L., Shcheglovitov, A., Portmann, T., Li, Y., Lee-Messer, C., Dolmetsch, R.E., Tsien, R.W., and Crabtree, G.R. (2011). MicroRNA-mediated conversion of human fibroblasts to neurons. Nature 476, 228–231. 10.1038/nature10323 [doi].

18. Wu, J.I., Lessard, J., Olave, I.A., Qiu, Z., Ghosh, A., Graef, I.A., and Crabtree, G.R. (2007). Regulation of dendritic development by neuron-specific chromatin remodeling complexes. Neuron 56, 94–108. S0896-6273(07)00664-2 [pii].

19. Vogel-Ciernia, A., Matheos, D.P., Barrett, R.M., Kramar, E.A., Azzawi, S., Chen, Y., Magnan, C.N., Zeller, M., Sylvain, A., Haettig, J., et al. (2013). The neuron-specific chromatin regulatory subunit BAF53b is necessary for synaptic plasticity and memory. Nat. Neurosci. 16, 552–561. 10.1038/nn.3359 [doi].

20. Tuoc, T.C., Boretius, S., Sansom, S.N., Pitulescu, M.E., Frahm, J., Livesey, F.J., and Stoykova, A. (2013). Chromatin regulation by BAF170 controls cerebral cortical size and thickness. Dev. Cell 25, 256–269. 10.1016/j.devcel.2013.04.005 [doi].

21. Matsumoto, S., Banine, F., Struve, J., Xing, R., Adams, C., Liu, Y., Metzger, D., Chambon, P., Rao, M.S., and Sherman, L.S. (2006). Brg1 is required for murine neural stem cell maintenance and gliogenesis. Dev. Biol. 289, 372–383. S0012-1606(05)00770-0 [pii].

22. Reyes, J.C., Barra, J., Muchardt, C., Camus, A., Babinet, C., and Yaniv, M. (1998). Altered control of cellular proliferation in the absence of mammalian brahma (SNF2alpha). EMBO J. 17, 6979– 6991. 10.1093/emboj/17.23.6979 [doi].

23. Hoyer, J., Ekici, A.B., Endele, S., Popp, B., Zweier, C., Wiesener, A., Wohlleber, E., Dufke, A., Rossier, E., Petsch, C., et al. (2012). Haploinsufficiency of ARID1B, a member of the SWI/SNF-a chromatin-remodeling complex, is a frequent cause of intellectual disability. Am. J. Hum. Genet. 90, 565–572. 10.1016/j.ajhg.2012.02.007 [doi].

24. Halgren, C., Kjaergaard, S., Bak, M., Hansen, C., El-Schich, Z., Anderson, C.M., Henriksen, K.F., Hjalgrim, H., Kirchhoff, M., Bijlsma, E.K., et al. (2012). Corpus callosum abnormalities, intellectual disability, speech impairment, and autism in patients with haploinsufficiency of ARID1B. Clin. Genet. 82, 248–255. 10.1111/j.1399-0004.2011.01755.x [doi].

25. Study, D.D.D. (2015). Large-scale discovery of novel genetic causes of developmental disorders. Nature 519, 223–228. 10.1038/nature14135 [doi].

26. Santen, G.W., Aten, E., Sun, Y., Almomani, R., Gilissen, C., Nielsen, M., Kant, S.G., Snoeck, I.N., Peeters, E.A., Hilhorst-Hofstee, Y., et al. (2012). Mutations in SWI/SNF chromatin remodeling complex gene ARID1B cause Coffin-Siris syndrome. Nat. Genet. 44, 379–380. 10.1038/ng.2217 [doi].

27. Zhang, Z., Cao, M., Chang, C.W., Wang, C., Shi, X., Zhan, X., Birnbaum, S.G., Bezprozvanny, I., Huber, K.M., and Wu, J.I. (2015). Autism-Associated Chromatin Regulator Brg1/SmarcA4 Is Required for Synapse Development and Myocyte Enhancer Factor 2-Mediated Synapse Remodeling. Mol. Cell. Biol. 36, 70–83. 10.1128/MCB.00534-15 [doi].

28. Valencia, A.M., Sankar, A., van der Sluijs, P.J., Satterstrom, F.K., Fu, J., Talkowski, M.E., Vergano, S.A.S., Santen, G.W.E., and Kadoch, C. (2023). Landscape of mSWI/SNF chromatin remodeling complex perturbations in neurodevelopmental disorders. Nat. Genet. 10.1038/s41588-023-01451-6.

29. Alfert, A., Moreno, N., and Kerl, K. (2019). The BAF complex in development and disease. Epigenetics and Chromatin. 10.1186/s13072-019-0264-y.

30. Peterson, C.L., and Herskowitz, I. (1992). Characterization of the yeast SWI1, SWI2, and SWI3 genes, which encode a global activator of transcription. Cell 68, 573–583. 10.1016/0092-8674(92)90192-F.

31. Hirschhorn, J.N., Brown, S.A., Clark, C.D., and Winston, F. (1992). Evidence that SNF2/SWI2 and SNF5 activate transcription in yeast by altering chromatin structure. Genes Dev. 6, 2288–2298. 10.1101/gad.6.12a.2288.

32. Păun, O., Tan, Y.X., Patel, H., Strohbuecker, S., Ghanate, A., Cobolli-Gigli, C., Llorian Sopena, M., Gerontogianni, L., Goldstone, R., Ang, S.-L., et al. (2023). Pioneer factor ASCL1 cooperates with the mSWI/SNF complex at distal regulatory elements to regulate human neural differentiation. Genes Dev. 37, 218–242. 10.1101/gad.350269.122.

33. Alver, B.H., Kim, K.H., Lu, P., Wang, X., Manchester, H.E., Wang, W., Haswell, J.R., Park, P.J., and Roberts, C.W.M. (2017). The SWI/SNF chromatin remodelling complex is required for maintenance of lineage specific enhancers. Nat. Commun. 8, 14648. 10.1038/ncomms14648.

34. Mathur, R., Alver, B.H., San Roman, A.K., Wilson, B.G., Wang, X., Agoston, A.T., Park, P.J., Shivdasani, R.A., and Roberts, C.W.M. (2017). ARID1A loss impairs enhancer-mediated gene regulation and drives colon cancer in mice. Nat. Genet. 49, 296–302. 10.1038/ng.3744.

35. Nakayama, R.T., Pulice, J.L., Valencia, A.M., McBride, M.J., McKenzie, Z.M., Gillespie, M.A., Ku, W.L., Teng, M., Cui, K., Williams, R.T., et al. (2017). SMARCB1 is required for widespread BAF complex–mediated activation of enhancers and bivalent promoters. Nat. Genet. 49, 1613–1623. 10.1038/ng.3958.

36. Park, Y.-K., Lee, J.-E., Yan, Z., McKernan, K., O’Haren, T., Wang, W., Peng, W., and Ge, K. (2021). Interplay of BAF and MLL4 promotes cell type-specific enhancer activation. Nat. Commun. 12, 1630. 10.1038/s41467-021-21893-y.

37. Brahma, S., and Henikoff, S. (2023). The BAF chromatin remodeler synergizes with RNA polymerase II and transcription factors to evict nucleosomes. Nat. Genet. 10.1038/s41588-023-01603-8.

38. Oruba, A., Saccani, S., and van Essen, D. (2020). Role of cell-type specific nucleosome positioning in inducible activation of mammalian promoters. Nat. Commun. 11, 1075. 10.1038/s41467-020-14950-5.

39. Fujinaga, K., Huang, F., and Peterlin, B.M. (2023). P-TEFb: The master regulator of transcription elongation. Mol. Cell 83, 393–403. 10.1016/j.molcel.2022.12.006.

40. Cramer, P. (2019). Eukaryotic Transcription Turns 50. Cell 179, 808–812. 10.1016/j.cell.2019.09.018.

41. Mohamed, A.A., Vazquez Nunez, R., and Vos, S.M. (2022). Structural advances in transcription elongation. Curr. Opin. Struct. Biol. 75, 102422. 10.1016/j.sbi.2022.102422.

42. Vos, S.M., Farnung, L., Linden, A., Urlaub, H., and Cramer, P. (2020). Structure of complete Pol II–DSIF–PAF–SPT6 transcription complex reveals RTF1 allosteric activation. Nat. Struct. Mol. Biol. 27, 668–677. 10.1038/s41594-020-0437-1.

43. Kulaeva, O.I., Hsieh, F.-K., and Studitsky, V.M. (2010). RNA polymerase complexes cooperate to relieve the nucleosomal barrier and evict histones. Proc. Natl. Acad. Sci. 107, 11325–11330. 10.1073/pnas.1001148107.

44. A., S.M., and Kevin, S. (2007). The Swi/Snf Complex Is Important for Histone Eviction during Transcriptional Activation and RNA Polymerase II Elongation In Vivo. Mol. Cell. Biol. 27, 6987– 6995. 10.1128/MCB.00717-07.

45. Subtil-Rodríguez, A., and Reyes, J.C. (2010). BRG1 helps RNA polymerase II to overcome a nucleosomal barrier during elongation, in vivo. EMBO Rep. 11, 751–757. 10.1038/embor.2010.131.

46. Armstrong, J.A., Papoulas, O., Daubresse, G., Sperling, A.S., Lis, J.T., Scott, M.P., and Tamkun, J.W. (2002). The Drosophila BRM complex facilitates global transcription by RNA polymerase II. EMBO J. 21, 5245–5254. 10.1093/emboj/cdf517.

47. Batsché, E., Yaniv, M., and Muchardt, C. (2006). The human SWI/SNF subunit Brm is a regulator of alternative splicing. Nat. Struct. Mol. Biol. 13, 22–29. 10.1038/nsmb1030.

48. Corey, L.L., Weirich, C.S., Benjamin, I.J., and Kingston, R.E. (2003). Localized recruitment of a chromatin-remodeling activity by an activator in vivo drives transcriptional elongation. Genes Dev. 17, 1392–1401. 10.1101/gad.1071803.

49. Mazina, M.Y., Nikolenko, Y. V, Krasnov, A.N., and Vorobyeva, N.E. (2016). SWI/SNF protein complexes participate in the initiation and elongation stages of Drosophila hsp70 gene transcription. Russ. J. Genet. 52, 141–145. 10.1134/S1022795416010105.

50. Tyssowski, K.M., DeStefino, N.R., Cho, J.-H., Dunn, C.J., Poston, R.G., Carty, C.E., Jones, R.D., Chang, S.M., Romeo, P., Wurzelmann, M.K., et al. (2018). Different Neuronal Activity Patterns Induce Different Gene Expression Programs. Neuron 98, 530–546.e11. 10.1016/j.neuron.2018.04.001.

51. Saha, R.N., Wissink, E.M., Bailey, E.R., Zhao, M., Fargo, D.C., Hwang, J.-Y., Daigle, K.R., Fenn, J.D., Adelman, K., and Dudek, S.M. (2011). Rapid activity-induced transcription of Arc and other IEGs relies on poised RNA polymerase II. Nat. Neurosci. 14. 10.1038/nn.2839.

52. Fujita, T., Ryser, S., Piuz, I., and Schlegel, W. (2008). Up-Regulation of P-TEFb by the MEK1-Extracellular Signal-Regulated Kinase Signaling Pathway Contributes to Stimulated Transcription Elongation of Immediate Early Genes in Neuroendocrine Cells. Mol. Cell. Biol. 28, 1630–1643. 10.1128/MCB.01767-07.

53. Farnaby, W., Koegl, M., Roy, M.J., Whitworth, C., Diers, E., Trainor, N., Zollman, D., Steurer, S., Karolyi-Oezguer, J., Riedmueller, C., et al. (2019). BAF complex vulnerabilities in cancer demonstrated via structure-based PROTAC design. Nat. Chem. Biol. 10.1038/s41589-019-0294-6.

54. Dunn, C.J., Sarkar, P., Bailey, E.R., Farris, S., Zhao, M., Ward, J.M., Dudek, S.M., and Saha, R.N. (2017). Histone hypervariants H2A.Z.1 and H2A.Z.2 play independent and context-specific roles in neuronal activity-induced transcription of Arc/ Arg3.1 and other immediate early genes. eNeuro. 10.1523/ENEURO.0040-17.2017.

55. Rienecker, K.D.A., Poston, R.G., Segales, J.S., Finholm, I.W., Sono, M.H., Munteanu, S.J., Ghaninejad-Esfahani, M., Rejepova, A., Tejeda-Garibay, S., Wickman, K., et al. (2022). Mild membrane depolarization in neurons induces immediate early gene transcription and acutely subdues responses to successive stimulus. J. Biol. Chem., 102278. 10.1016/J.JBC.2022.102278.

56. Papillon, J.P.N., Nakajima, K., Adair, C.D., Hempel, J., Jouk, A.O., Karki, R.G., Mathieu, S., Möbitz, H., Ntaganda, R., Smith, T., et al. (2018). Discovery of Orally Active Inhibitors of Brahma Homolog (BRM)/SMARCA2 ATPase Activity for the Treatment of Brahma Related Gene 1 (BRG1)/SMARCA4-Mutant Cancers. J. Med. Chem. 10.1021/acs.jmedchem.8b01318.

57. Battistello, E., Hixon, K.A., Comstock, D.E., Collings, C.K., Chen, X., Rodriguez Hernaez, J., Lee, S., Cervantes, K.S., Hinkley, M.M., Ntatsoulis, K., et al. (2023). Stepwise activities of mSWI/SNF family chromatin remodeling complexes direct T cell activation and exhaustion. Mol. Cell 83, 1216–1236.e12. 10.1016/j.molcel.2023.02.026.

58. Poston, R.G., Dunn, C.J., Sarkar, P., and Saha, R.N. (2018). Persistent 6-OH-BDE-47 exposure impairs functional neuronal maturation and alters expression of neurodevelopmentally-relevant chromatin remodelers. Environ. Epigenetics 4, 1–15. 10.1093/eep/dvx020.

59. Marian, C.A., Stoszko, M., Wang, L., Leighty, M.W., de Crignis, E., Maschinot, C.A., Gatchalian, J., Carter, B.C., Chowdhury, B., Hargreaves, D.C., et al. (2018). Small Molecule Targeting of Specific BAF (mSWI/SNF) Complexes for HIV Latency Reversal. Cell Chem. Biol. 10.1016/j.chembiol.2018.08.004.

60. Jafari, R., Almqvist, H., Axelsson, H., Ignatushchenko, M., Lundbäck, T., Nordlund, P., and Molina, D.M. (2014). The cellular thermal shift assay for evaluating drug target interactions in cells. Nat. Protoc. 9, 2100–2122. 10.1038/nprot.2014.138.

61. Ordonez-Rubiano, S.C., Maschinot, C.A., Wang, S., Sood, S., Baracaldo-Lancheros, L.F., Strohmier, B.P., McQuade, A.J., Smith, B.C., and Dykhuizen, E.C. (2023). Rational Design and Development of Selective BRD7 Bromodomain Inhibitors and Their Activity in Prostate Cancer. J. Med. Chem. 66, 11250–11270. 10.1021/acs.jmedchem.3c00671.

62. Theodoulou, N.H., Bamborough, P., Bannister, A.J., Becher, I., Bit, R.A., Che, K.H., Chung, C., Dittmann, A., Drewes, G., Drewry, D.H., et al. (2016). Discovery of I-BRD9, a Selective Cell Active Chemical Probe for Bromodomain Containing Protein 9 Inhibition. J. Med. Chem. 59, 1425–1439. 10.1021/acs.jmedchem.5b00256.

63. Zoppi, V., Hughes, S.J., Maniaci, C., Testa, A., Gmaschitz, T., Wieshofer, C., Koegl, M., Riching, K.M., Daniels, D.L., Spallarossa, A., et al. (2019). Iterative Design and Optimization of Initially Inactive Proteolysis Targeting Chimeras (PROTACs) Identify VZ185 as a Potent, Fast, and Selective von Hippel-Lindau (VHL) Based Dual Degrader Probe of BRD9 and BRD7. J. Med. Chem. 10.1021/acs.jmedchem.8b01413.

64. Ma, H., Khaled, H.G., Wang, X., Mandelberg, N.J., Cohen, S.M., He, X., and Tsien, R.W. (2023). Excitation–transcription coupling, neuronal gene expression and synaptic plasticity. Nat. Rev. Neurosci. 24, 672–692. 10.1038/s41583-023-00742-5.

65. Nechaev, S., and Adelman, K. (2008). Promoter-proximal Pol II: When stalling speeds things up. Cell Cycle 7, 1539–1544. 10.4161/cc.7.11.6006.

66. Aoi, Y., Takahashi, Y., Shah, A.P., Iwanaszko, M., Rendleman, E.J., Khan, N.H., Cho, B.-K., Goo, Y.A., Ganesan, S., Kelleher, N.L., et al. (2021). SPT5 stabilization of promoter-proximal RNA polymerase II. Mol. Cell 81, 4413–4424.e5. 10.1016/j.molcel.2021.08.006.

67. Madabhushi, R., and Kim, T.-K. (2018). Emerging themes in neuronal activity-dependent gene expression. Mol. Cell. Neurosci. 87, 27–34. 10.1016/j.mcn.2017.11.009.

68. Peterlin, B.M., and Price, D.H. (2006). Controlling the Elongation Phase of Transcription with P-TEFb. Mol. Cell 23, 297–305. 10.1016/j.molcel.2006.06.014.

69. Vos, S.M., Farnung, L., Boehning, M., Wigge, C., Linden, A., Urlaub, H., and Cramer, P. (2018). Structure of activated transcription complex Pol II–DSIF–PAF–SPT6. Nature 560, 607–612. 10.1038/s41586-018-0440-4.

70. Olson, C.M., Jiang, B., Erb, M.A., Liang, Y., Doctor, Z.M., Zhang, Z., Zhang, T., Kwiatkowski, N., Boukhali, M., Green, J.L., et al. (2018). Pharmacological perturbation of CDK9 using selective CDK9 inhibition or degradation. Nat. Chem. Biol. 14, 163–170. 10.1038/nchembio.2538.

71. Ramakrishnan, R., and Rice, A.P. (2012). Cdk9 T-loop phosphorylation is regulated by the calcium signaling pathway. J. Cell. Physiol. 227, 609–617. 10.1002/jcp.22760.

72. Jonkers, I., Kwak, H., and Lis, J.T. (2014). Genome-wide dynamics of Pol II elongation and its interplay with promoter proximal pausing, chromatin, and exons. Elife 3, e02407. 10.7554/eLife.02407.

73. Zhang, H., Pandey, S., Travers, M., Sun, H., Morton, G., Madzo, J., Chung, W., Khowsathit, J., Perez-Leal, O., Barrero, C.A., et al. (2018). Targeting CDK9 Reactivates Epigenetically Silenced Genes in Cancer. Cell 175, 1244–1258.e26. 10.1016/j.cell.2018.09.051.

74. Aoi, Y., and Shilatifard, A. (2023). Transcriptional elongation control in developmental gene expression, aging, and disease. Mol. Cell 83, 3972–3999. 10.1016/j.molcel.2023.10.004.

75. Chen, F., Gao, X., and Shilatifard, A. (2015). Stably paused genes revealed through inhibition of transcription initiation by the TFIIH inhibitor triptolide. Genes Dev. 29, 39–47. 10.1101/gad.246173.114.

76. Elrod, N.D., Henriques, T., Huang, K.-L., Tatomer, D.C., Wilusz, J.E., Wagner, E.J., and Adelman, K. (2019). The Integrator Complex Attenuates Promoter-Proximal Transcription at Protein-Coding Genes. Mol. Cell 76, 738–752.e7. 10.1016/j.molcel.2019.10.034.

77. Gourisankar, S., Wenderski, W., Paulo, J.A., Kim, S.H., Roepke, K., Ellis, C., Gygi, S.P., and Crabtree, G.R. (2023). Synaptic Activity Causes Minute-scale Changes in BAF Complex Composition and Function. bioRxiv, 2023.10.13.562244. 10.1101/2023.10.13.562244.

78. Iurlaro, M., Stadler, M.B., Masoni, F., Jagani, Z., Galli, G.G., and Schübeler, D. (2021). Mammalian SWI/SNF continuously restores local accessibility to chromatin. Nat. Genet. 10.1038/s41588-020-00768-w.

79. Hoffman, J.A., Trotter, K.W., Ward, J.M., and Archer, T.K. (2018). BRG1 governs glucocorticoid receptor interactions with chromatin and pioneer factors across the genome. Elife 7, e35073. 10.7554/eLife.35073.

80. Kim, B., Luo, Y., Zhan, X., Zhang, Z., Shi, X., Yi, J., Xuan, Z., and Wu, J. (2021). Neuronal activity-induced BRG1 phosphorylation regulates enhancer activation. Cell Rep. 36. 10.1016/j.celrep.2021.109357.

81. Zhang, X., Li, B., Li, W., Ma, L., Zheng, D., Li, L., Yang, W., Chu, M., Chen, W., Mailman, R.B., et al. (2014). Transcriptional Repression by the BRG1-SWI/SNF Complex Affects the Pluripotency of Human Embryonic Stem Cells. Stem Cell Reports 3, 460–474. 10.1016/j.stemcr.2014.07.004.

82. Tolstorukov, M.Y., Sansam, C.G., Lu, P., Koellhoffer, E.C., Helming, K.C., Alver, B.H., Tillman, E.J., Evans, J.A., Wilson, B.G., Park, P.J., et al. (2013). Swi/Snf chromatin remodeling/tumor suppressor complex establishes nucleosome occupancy at target promoters. Proc. Natl. Acad. Sci. 110, 10165–10170. 10.1073/pnas.1302209110.

83. Trizzino, M., Barbieri, E., Petracovici, A., Wu, S., Welsh, S.A., Owens, T.A., Licciulli, S., Zhang, R., and Gardini, A. (2018). The Tumor Suppressor ARID1A Controls Global Transcription via Pausing of RNA Polymerase II. Cell Rep. 10.1016/j.celrep.2018.05.097.

84. Mayer, A., di Iulio, J., Maleri, S., Eser, U., Vierstra, J., Reynolds, A., Sandstrom, R., Stamatoyannopoulos, J.A., and Churchman, L.S. (2015). Native Elongating Transcript Sequencing Reveals Human Transcriptional Activity at Nucleotide Resolution. Cell 161, 541–554. 10.1016/j.cell.2015.03.010.

85. Wenderski, W., Wang, L., Krokhotin, A., Walsh, J.J., Li, H., Shoji, H., Ghosh, S., George, R.D., Miller, E.L., Elias, L., et al. (2020). Loss of the neural-specific BAF subunit ACTL6B relieves repression of early response genes and causes recessive autism. Proc. Natl. Acad. Sci. U. S. A. 10.1073/pnas.1908238117.

86. Modur, V., Singh, N., Mohanty, V., Chung, E., Muhammad, B., Choi, K., Chen, X., Chetal, K., Ratner, N., Salomonis, N., et al. (2018). Defective transcription elongation in a subset of cancers confers immunotherapy resistance. Nat. Commun. 9, 4410. 10.1038/s41467-018-06810-0.

87. Miller, T.E., Liau, B.B., Wallace, L.C., Morton, A.R., Xie, Q., Dixit, D., Factor, D.C., Kim, L.J.Y., Morrow, J.J., Wu, Q., et al. (2017). Transcription elongation factors represent in vivo cancer dependencies in glioblastoma. Nature 547, 355–359. 10.1038/nature23000.

88. Muhammad, B., Parks, L.G., Komurov, K., and Privette Vinnedge, L.M. (2022). Defective transcription elongation in human cancers imposes targetable proteotoxic vulnerability. Transl. Oncol. 16, 101323. 10.1016/j.tranon.2021.101323.

89. St. Pierre, R., and Kadoch, C. (2017). Mammalian SWI/SNF complexes in cancer: emerging therapeutic opportunities. Curr. Opin. Genet. Dev. 42, 56–67. 10.1016/j.gde.2017.02.004.

90. King, I.F., Yandava, C.N., Mabb, A.M., Hsiao, J.S., Huang, H.S., Pearson, B.L., Calabrese, J.M., Starmer, J., Parker, J.S., Magnuson, T., et al. (2013). Topoisomerases facilitate transcription of long genes linked to autism. Nature 501, 58–62. 10.1038/nature12504.

91. Qiu, Z., Zhao, L., Shen, J.Z., Liang, Z., Wu, Q., Yang, K., Min, L., Gimple, R.C., Yang, Q., Bhargava, S., et al. (2022). Transcription Elongation Machinery Is a Druggable Dependency and Potentiates Immunotherapy in Glioblastoma Stem Cells. Cancer Discov. 12, 502–521. 10.1158/2159-8290.CD-20-1848.

92. Bidart, M., El Atifi, M., Miladi, S., Rendu, J., Satre, V., Ray, P.F., Bosson, C., Devillard, F., Lehalle, D., Malan, V., et al. (2017). Microduplication of the ARID1A gene causes intellectual disability with recognizable syndromic features. Genet. Med. 10.1038/gim.2016.180.

93. Machol, K., Rousseau, J., Ehresmann, S., Garcia, T., Nguyen, T.T.M., Spillmann, R.C., Sullivan, J.A., Shashi, V., Jiang, Y. hui, Stong, N., et al. (2019). Expanding the Spectrum of BAF-Related Disorders: De Novo Variants in SMARCC2 Cause a Syndrome with Intellectual Disability and Developmental Delay. Am. J. Hum. Genet. 10.1016/j.ajhg.2018.11.007.

94. Neale, B.M., Kou, Y., Liu, L., Ma’ayan, A., Samocha, K.E., Sabo, A., Lin, C.F., Stevens, C., Wang, L.S., Makarov, V., et al. (2012). Patterns and rates of exonic de novo mutations in autism spectrum disorders. Nature 485, 242–245. 10.1038/nature11011 [doi].

95. Iossifov, I., Levy, D., Allen, J., Ye, K., Ronemus, M., Lee, Y.H., Yamrom, B., and Wigler, M. (2015). Low load for disruptive mutations in autism genes and their biased transmission. Proc. Natl. Acad. Sci. U. S. A. 10.1073/pnas.1516376112.

96. Takata, A., Ionita-Laza, I., Gogos, J.A., Xu, B., and Karayiorgou, M. (2016). De Novo Synonymous Mutations in Regulatory Elements Contribute to the Genetic Etiology of Autism and Schizophrenia. Neuron. 10.1016/j.neuron.2016.02.024.

97. Bosch, E., Popp, B., Güse, E., Skinner, C., van der Sluijs, P.J., Maystadt, I., Pinto, A.M., Renieri, A., Bruno, L.P., Granata, S., et al. (2023). Elucidating the clinical and molecular spectrum of SMARCC2-associated NDD in a cohort of 65 affected individuals. Genet. Med. 25, 100950. 10.1016/j.gim.2023.100950.

98. Navickas, S.M., Giles, K.A., Brettingham-Moore, K.H., and Taberlay, P.C. (2023). The role of chromatin remodeler SMARCA4/BRG1 in brain cancers: a potential therapeutic target. Oncogene 42, 2363–2373. 10.1038/s41388-023-02773-9.

99. Dunn, C.J., Sarkar, P., Bailey, E.R., Farris, S., Zhao, M., Ward, J.M., Dudek, S.M., and Saha, R.N. (2017). Histone hypervariants H2A.Z.1 and H2A.Z.2 play independent and context-specific roles in neuronal activity-induced transcription of Arc/ Arg3.1 and other immediate early genes. eNeuro 4. 10.1523/ENEURO.0040-17.2017.

