## Supplemental Figures (S1-S3) for "Activity-assembled nBAF complex mediates rapid immediate early gene transcription by regulating RNA Polymerase II productive elongation"

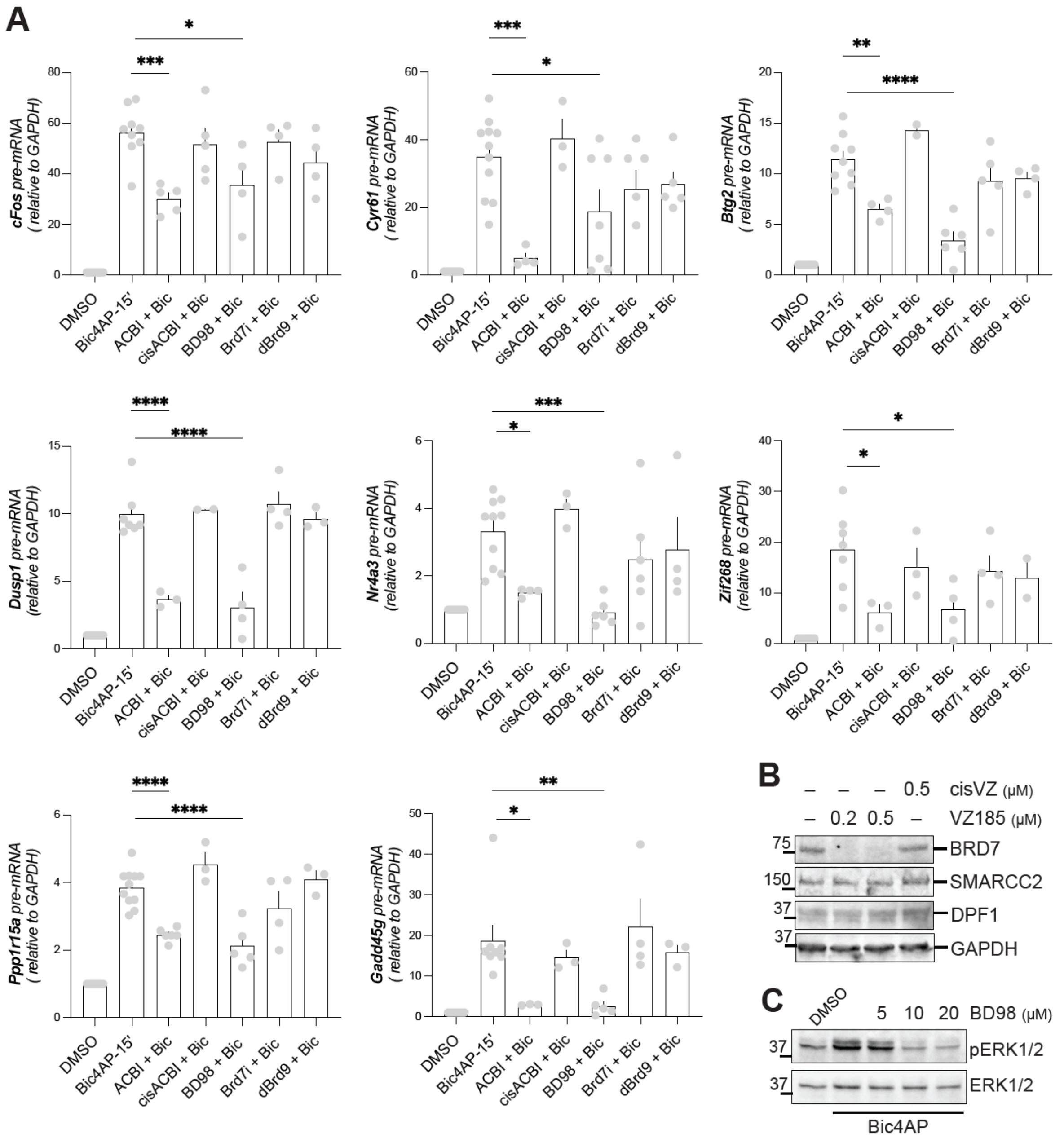


Cornejo *et. al.* figure S1

Figure S1: **nBAF complex is required for optimal transcription of rIEGs.**
A) Multiple rIEGs pre-mRNA levels normalized to GAPDH. Neurons were treated with ACBI1, cis-ACBI1, BD98, BRD7i, Brd9i, which was followed by induction of neuronal activity via Bic+4AP treatment for 15 minutes. Gene names are indicated on the Y-axis. Grey dots represent biological replicates; errors bars show SE of the mean. **P*<0.05; ***P*<0.01. One-way ANOVAs were performed followed by Tukey's multiple comparisons test as *post-hoc.* B) Representative western blots of whole-cell lysates to show the effect of VZ185, which degrades BRD7, not nBAF subunits. Neurons were treated with VZ185 or cisVZ for three hours at indicated concentrations. C) Neurons were treated with DMSO (control), 5μM, 10μM, or 20μM BD98, followed by 10 minutes of Bic+4AP. Blot was probed for total ERK1/2 and phosho-ERK1/2.

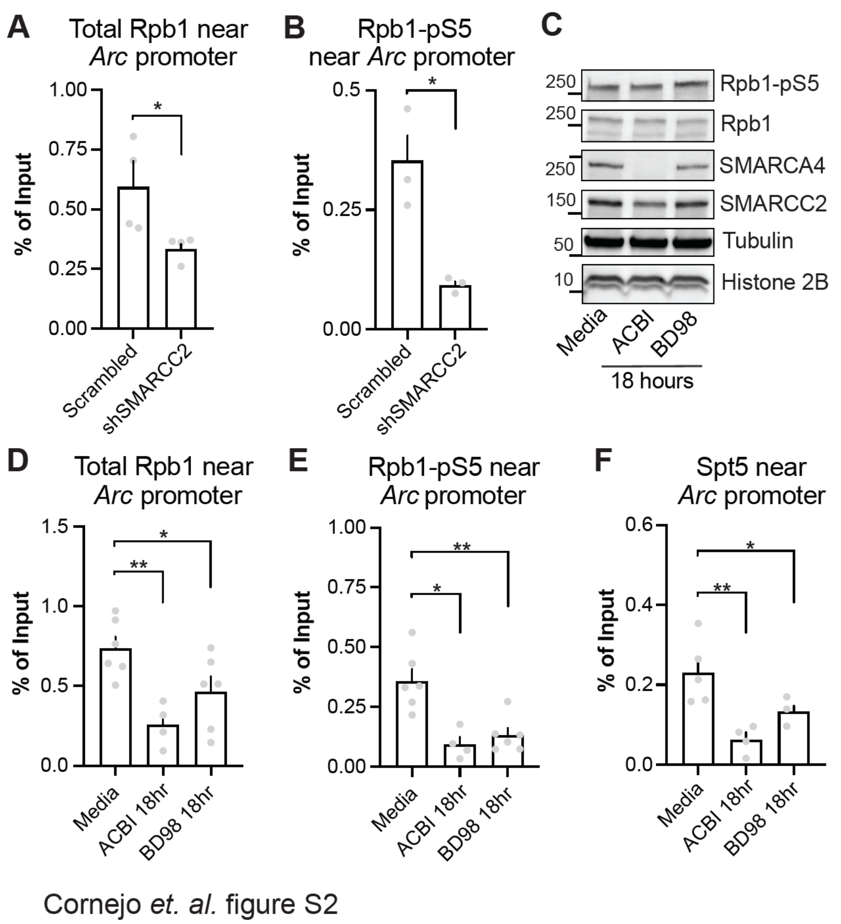


Figure S2: **Prolonged degradation, inhibition, or depletion of BAF complex results in attenuated promoter-proximal RNA Pol II pausing near *Arc* promoter.**

A and B) Neurons were infected with lentiviruses to deliver shRNA against SMARCC2 or control (Scrambled). Quantified total RNA Pol2 binding near *Arc* promoter was determined by ChIP with antibody against Rpb1-NTD and paused RNA Pol2 with Rpb1-pS5. C) Neurons were treated with ACBI1 (2.5 µM) or BD98 (5 µM) for 18 hours. Whole cell lysates were then analyzed via western blot to assess protein levels of RNA Pol II or BAF subunits. D, E and F) ChIP data depicting SMARCC2 binding near *Arc* promoter after 18-hour treatment with ACBI or BD98. ChIP was performed for total RNA Pol2, paused RNA Pol2 (Rpb1-pS5), and SPT5. For figures A-B, unpaired two- tailed t-test were performed. For figures, D-F, one-way ANOVAs were performed. Grey dots represent biological replicates; errors bars show SE of the mean. *$P<0.05$; **$P<0.01$.

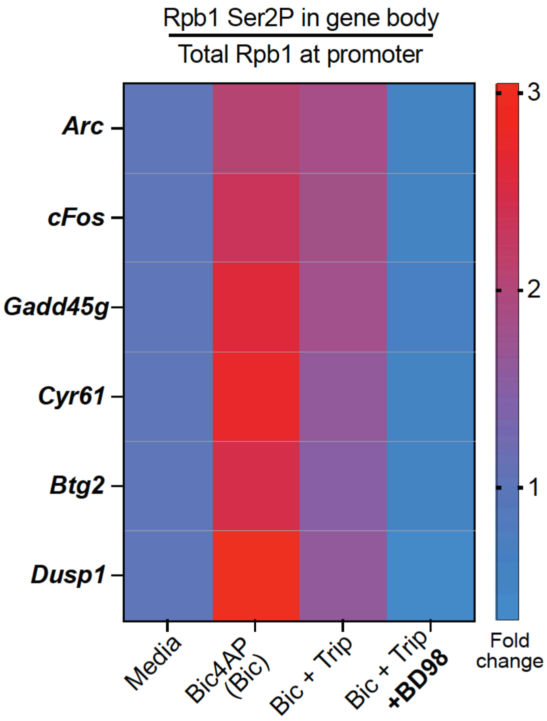


Cornejo *et. al.* figure S3

**Figure S3. The nBAF complex is required for Pol2 productive elongation in multiple rIEGs.** Heat map was generated using ChIP material described in figure 7C-7E. The heat map represents as fold change the ratio of elongation competent Pol2 in gene bodies normalized by total amount of Pol2 near corresponding promoters. Values for *Arc* are depicted here for cross reference with the main figure. qPCR with primers against promoters and gene body of five rIEGS were used to create additional rows in the heatmap. Heatmap represents the mean value of *N*=3. Bic+Trip+BD98 values were significantly less than Bic+Trip values in all cases.
